# A peptide-based screen for cell death inhibitors identifies the cytoprotective compound CDL36

**DOI:** 10.64898/2026.07.09.737573

**Authors:** Zintis Inde, Sebastian Keppler, Jesse Gelles, Cameron S. Fraser, Adam Presser, Jarvier N. Mohammed, May Jung, David S. Garvey, Tudor Moldoveanu, Jerry E. Chipuk, Kristopher A. Sarosiek

**Affiliations:** John B. Little Center for Radiation Sciences, Harvard T.H. Chan School of Public Health, Boston, MA; Laboratory of Systems Pharmacology, Harvard Program in Therapeutic Science, Department of Systems Biology, Harvard Medical School, Boston, MA; Department of Systems Biology, University of Massachusetts Chan Medical School, Worcester, MA; Departments of Oncological Sciences and Dermatology, Icahn School of Medicine at Mount Sinai, New York, NY; DSG Pharma Consulting LLC, Dover, MA; Department of Biochemistry and Molecular Biology, University of Arkansas for Medical Sciences, Little Rock, AR; Department of Medical Oncology, Dana-Farber Cancer Institute, Boston MA

## Abstract

Small molecule inhibitors of cell death have wide-ranging potential applications, both as tool compounds in the laboratory and as clinical modulators of pathologic cell death. Previous screening efforts have identified candidate compounds targeting the pro-apoptotic, pore-forming BCL-2 family proteins BAX and BAK, but the complex interactions of these proteins at the mitochondrial outer membrane (with other proteins and the membrane itself) present challenges for compound screening. Although no inhibitors of BAX or BAK have advanced to clinical testing to date, candidate inhibitors have thus far been identified via screening of membrane-containing systems such as liposomes and isolated mitochondria. To address some of the challenges of chemical screening for apoptosis inhibitors, we conducted a small molecule screen utilizing BH3 profiling, a method that quantifies mitochondrial outer membrane permeabilization (MOMP) upon treatment with pro-apoptotic peptides derived from BCL-2 family proteins. Of over 40,000 compounds screened, we identified a series of compounds that prevent MOMP in response to pro-apoptotic peptides. The most potent of these, CDL36, binds to BAX and prevents MOMP at early timepoints. In longer term viability assays, the cytoprotective effect of CDL36 is most potent against death induced by doxorubicin, a widely used chemotherapeutic agent that causes dose-limiting cardiovascular toxicity. Our results elucidate the mechanism of action of new and existing cell death inhibitors, providing a foundation for further development of these inhibitors and potential insights into the mechanisms mediating doxorubicin toxicity in patients.

## Introduction

Apoptosis is a highly regulated pathway by which cells die, both in normal physiology and in various pathological contexts^1^. The pharmacologic modulation of apoptosis has both enabled and benefitted from increasing insights into the molecular mechanisms of apoptotic cell death. The elucidation of the roles of pro- and anti-apoptotic B-cell lymphoma 2 (BCL-2) family proteins has enabled the development of molecules known as BCL-2 homology domain 3 (BH3) mimetics, which inhibit anti-apoptotic BCL-2 proteins to induce apoptosis and have found substantial utility as cancer therapeutics^2^. In turn, BH3 mimetics and other small molecules have enabled researchers to probe the fundamental biology of the BCL-2 family, gaining insight not only into cell death regulation, but related phenotypes including inflammation and senescence^3^.

While pro-apoptotic pharmacologic agents have proved useful in both the laboratory and the clinic, the development of anti-apoptotic small molecules presents novel challenges^4^. Inhibitors of caspases, the protease executioners of apoptotic cell death, are useful tools for the inhibition of apoptosis execution in the laboratory but have shown little success in clinical application, in part because they may lie too far downstream in the apoptotic signaling cascade to rescue mitochondrial fitness^5^. By contrast, the pro-apoptotic proteins BCL-2 associated X (BAX, encoded by *BAX*) and BCL-2 antagonist/killer 1 (BAK, encoded by *BAK1*) represent early, essential steps in the initiation of apoptosis. Upon activation, BAX and BAK proteins form pores in the outer mitochondrial membrane to allow the release of pro-apoptotic cytochrome *c* and other factors into the cytoplasm^6–8^. This mitochondrial outer membrane permeabilization (MOMP) drives the downstream activation of caspases and represents a key bottleneck in the apoptotic signaling pathway. Recent work has also demonstrated that even in the absence of caspase activation, MOMP can promote inflammatory signaling and other phenotypes including senescence^3,9,10^, highlighting the need for inhibitors acting at earlier stages of apoptotic signaling, both as tool compounds for mechanistic investigation and as candidates for clinical application.

*In vitro* chemical screens for inhibitors of BAX have previously been conducted, mainly relying on systems such as *in vitro* liposome assays to identify hits^11–13^, but these assays do not fully recapitulate the cellular milieu that is critical for proper regulation and function of BAX. *In silico* small molecule screens have also been conducted^14^, as well as computational and directed evolution efforts toward developing peptide inhibitors of BAX and BAK^15,16^. To our knowledge, however, no small molecule or peptide inhibitor has yet advanced to clinical testing. Here, we performed a small molecule screen in cells that have been gently permeabilized to identify candidate molecules that could prevent the induction of MOMP by BH3 peptides derived from pro-apoptotic BCL-2 family proteins^17,18^. These BH3 peptides can inhibit anti-apoptotic BCL-2 family proteins or directly activate BAX and BAK, providing a pro-apoptotic stimulus in a more complete cellular context^1^. Demonstrating the utility of this screening approach, we identify a series of novel inhibitors of MOMP, including small molecules with substantial structural similarity as well as others with highly distinct structures. We demonstrate that a subset of these inhibitors directly binds BAX, including the most potent compound, CDL36. Although this compound potently inhibits mitochondrial outer membrane permeabilization by BAX/BAK over short timescales, we find that long-term protection is specific to the DNA damaging agent doxorubicin. Our results provide insight into the mechanism of action of new and existing apoptosis-modulating small molecules, and CDL36 serves as a useful compound for further development of specific and potent inhibitors of death induced by doxorubicin and other cell death inducers.

## Results

### BH3 Profiling Screen Identifies Candidate Inhibitors of BAX and BAK

To identify potential small molecule inhibitors of the pore-forming proteins BAX and BAK, we performed a screen of 44,827 small molecules for their ability to preserve mitochondrial membrane potential in two apoptosis-sensitive human lymphoma cell lines, U937 and OCI-Ly1, treated with pro-apoptotic BH3 peptides (Figure 1A). Compounds screened included known bioactive molecules, including FDA approved drugs, and the ChemDiv 7 Targeted Diversity library, comprised of a structurally diverse set of compounds with drug-like properties and greater potential for protein binding (Table S1). This screening approach (Figure 1B) is based on the BH3 profiling assay, which has previously been demonstrated to quantify the likelihood of apoptosis initiation in clinical samples and thereby predict treatment response^19^. BH3 profiling utilizes peptides derived from the BCL-2 Homology 3 (BH3) domains of pro-apoptotic proteins, here including Bcl-2 Interacting Mediator of cell death (BIM, encoded by *BCL2L11*) and BH3 Interacting Domain death agonist (BID). In this assay, cells are gently permeabilized with digitonin to disrupt the plasma membrane and allow peptide entry while preserving mitochondrial membranes and other cellular components that may modulate apoptotic signaling. In typical BH3 profiling, a series of single peptides is used to quantify responses to a range of pro-apoptotic stimuli; for the primary screen, BIM and BID peptides, which preferentially activate BAX and BAK^20^, respectively, were used in combination to select for screening compounds with broad protective effects against both routes of MOMP activation. The response to pro-apoptotic peptide treatment was measured via JC-1 staining for mitochondrial membrane potential. Membrane potential is lost in a BAX/BAK-dependent manner upon BH3 peptide-induced mitochondrial outer membrane permeabilization (MOMP), and the loss of membrane potential causes the JC-1 dye to shift from red fluorescent aggregates to green fluorescent monomers^21^. Cell lines employed in the screen expressed both BAX and BAK (Figure 1C), allowing identification of compounds with potency against both pore-forming proteins. For each compound in each cell line, we calculated a per-plate z-score from the JC-1 fluorescence values obtained via plate reader assay (Figure 1D). For subsequent follow-up, we selected compounds with a z-score greater than 3 in both biological replicates in one or both cell lines, eliminating likely autofluorescent or BAX/BAK-independent compounds that gave a z-score greater than 1 in a separate screen of BH3 peptide-treated BAX/BAK double knockout HeLa cervical cancer cells.

**Figure 1.**
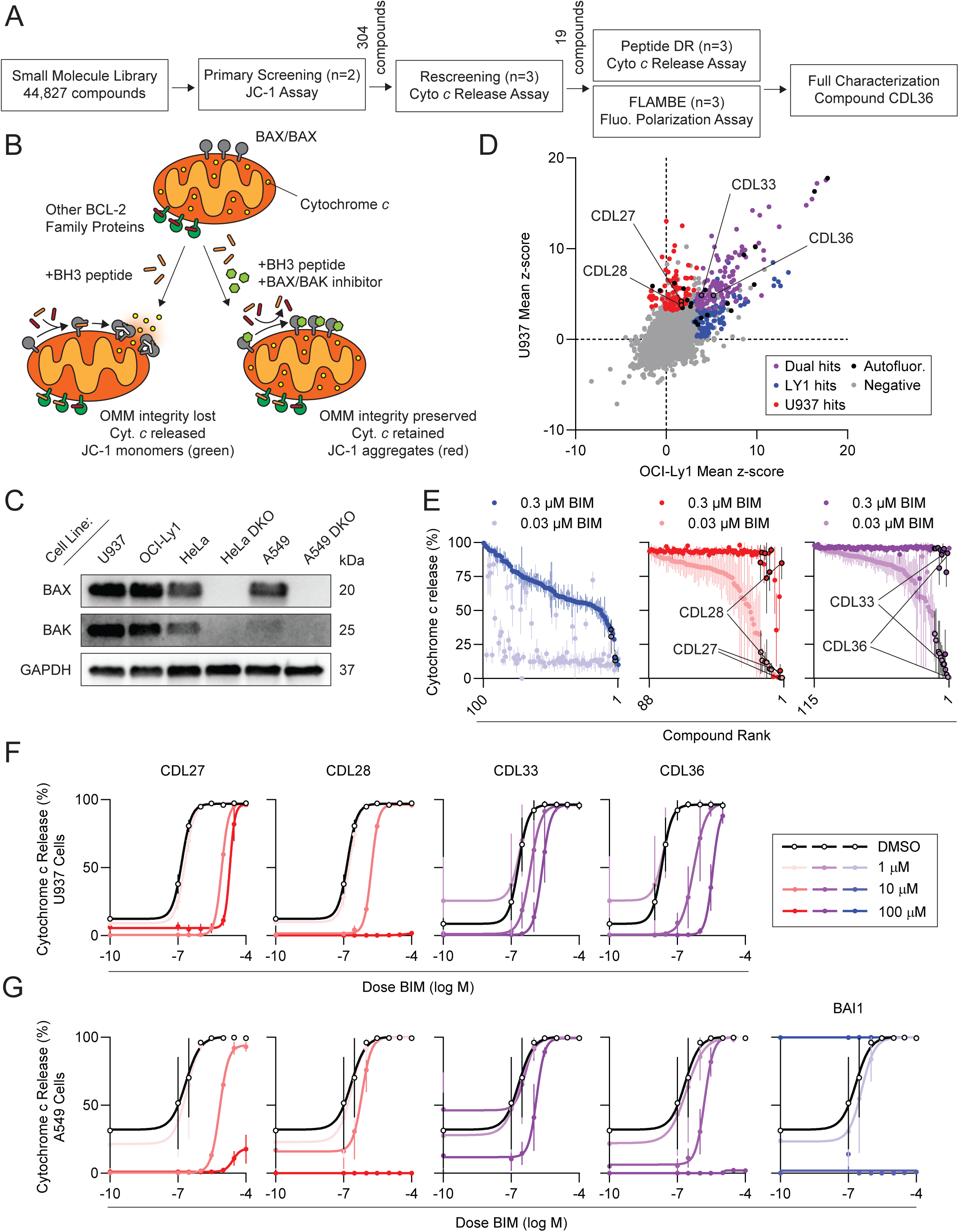
A BH3 profiling screen identifies inhibitors of mitochondrial outer membrane permeabilization. A. Schematic describing the screening and characterization of small molecules to identify inhibitors of mitochondrial outer membrane permeabilization (MOMP). B. Schematic depicting the expected effect of pro-apoptotic BH3 peptides on outer mitochondrial membrane integrity and apoptotic protein release in the presence or absence of a BAX/BAK inhibitor. C. Western blot analysis of cell lines used in primary screening and subsequent characterization. D. Activity of 44,827 small molecules in preventing MOMP as measured by JC-1 signal upon treatment with the combined pro-apoptotic peptides BIM and BID in U937 and OCI-LY1 cells. Each point represents the mean z-score of two technical replicates in each cell line. Hit compounds had a z-score ≥ 3 in both replicates in one or both cell lines, and autofluorescent compounds were removed based on a z-score ≥ 1 in a separate screen in BAX/BAK KO HeLa cells. E. Secondary screening of hit compounds in three libraries (as indicated in D) comprised of the compounds with activity in OCI-LY1, U937, or both cell lines. Data represent the mean and SD of cytochrome c signal as measured by flow cytometry upon treatment of OCI-LY1 cells (OCI-LY1 hit compounds, left) or U937 cells (U397 hit compounds, center; dual hit compounds, right) with BIM peptide at the indicated doses and candidate inhibitors at 100 µM. Data represent the mean and SD of % cytochrome c negativity as measured by flow cytometry. n = 3 biological replicates. F. Activity of selected hit compounds in U937 cells co-treated with the indicated compound doses and a range of BIM peptide doses. Data represent the mean and SD of % cytochrome c negativity. n = 3 biological replicates. G. Activity of selected hit compounds and reported BAX inhibitor BAI1 in A549 cells co-treated with the indicated compound doses and a range of BIM peptide doses. Data represent the mean and SD of % cytochrome c negativity. n = 3 biological replicates.

We screened the selected compounds in three sets of candidate inhibitors, comprised of compounds that were effective in the primary screen (z-score > 3) in OCI-Ly1 cells, U937 cells, or both (Figure 1E). We conducted secondary screening by an orthogonal, flow cytometry-based version of BH3 profiling using a fluorescent-conjugated antibody against cytochrome *c*. In this assay, fluorescent signal is lost (indicating cytochrome *c* release) upon peptide treatment at the indicated doses but retained in vehicle control treatments or conditions where the candidate inhibitor prevents MOMP. Candidate compounds identified as effective in U937 cells or in both cell lines were rescreened in U937 cells, whereas candidates identified in OCI-Ly1 cells were rescreened in OCI-Ly1 cells. Of the 35 candidate compounds that demonstrated the strongest protection against MOMP (greatest percentage retention of cytochrome *c* upon peptide treatment) in secondary screening, 19 were available for purchase from Chemdiv; we obtained these compounds and tested the efficacy of newly-obtained stocks (applied at 1, 10, and 100 µM doses) in preventing MOMP upon treatment with a range of BIM peptide doses in U937 cells (Figure 1F, S1A-B). The tested compounds showed a range of protective effects in these cells, with compounds from both the U937-specific and dual hit libraries showing decreased sensitivity to BIM (e.g. CDL27, CDL33, CDL36) or even complete protection against MOMP at the tested BIM doses (e.g. CDL28).

To test how generalizable the observed protective effects were, we further tested selected hit compounds in two adherent cell lines, HeLa and A549 lung adenocarcinoma, treated with the same range of BIM doses (Figure 1G, S1C). In these cells, we also tested the previously reported BAX inhibitor BAI1, allowing us to compare its protective effect against the novel candidate inhibitors. In both cell lines, BAI1 showed BIM-independent toxicity at the 100 µM dose, prompting us to use this compound at 10 µM for subsequent analyses. The two candidate compounds with the strongest protective effect in A549 cells, CDL28 and CDL36, also showed protection against peptide-induced MOMP in HeLa cells, and the effect of both compounds at a 100 µM dose was roughly comparable with the 10 µM dose of BAI1, consistent with the reported BAX-inhibitory effect of this compound. Thus, our screening pipeline identified molecules with potent anti-apoptotic effects across a range of cancer cell lines from diverse tissues of origin, but the mechanism for their activity was unknown. Therefore, we sought to determine which of these compounds bound directly to BAX or BAK.

### Screening Hits Directly Bind BAX in vitro

To test the binding of our candidate inhibitors to BAX and BAK, we first employed a fluorescence polarization assay: Fluorescence polarization Ligand Assay for Monitoring BAX Early-activation (FLAMBE)^22^. This *in vitro* assay utilizes a fluorophore-conjugated BAX-binding peptide whose polarization and depolarization indicates the binding and dissociation, respectively, of the peptide to and from recombinant BAX in solution^22^. We conducted FLAMBE analysis of the 19 candidate inhibitors at doses ranging from 39 nM to 5 µM. This analysis identified a limited set of compounds, including CDL36, whose addition to the peptide-BAX mixture prevented fluorescence polarization, thereby indicating a dose-dependent disruption of the binding between peptide and BAX (Figure 2A-C, S2A-B). By contrast, other compounds that had showed strong protection against apoptosis initiation (e.g. CDL28) did not substantially modulate BAX polarization kinetics, suggesting that their protective activity was unlikely to be mediated by inhibition of BAX.

**Figure 2.**
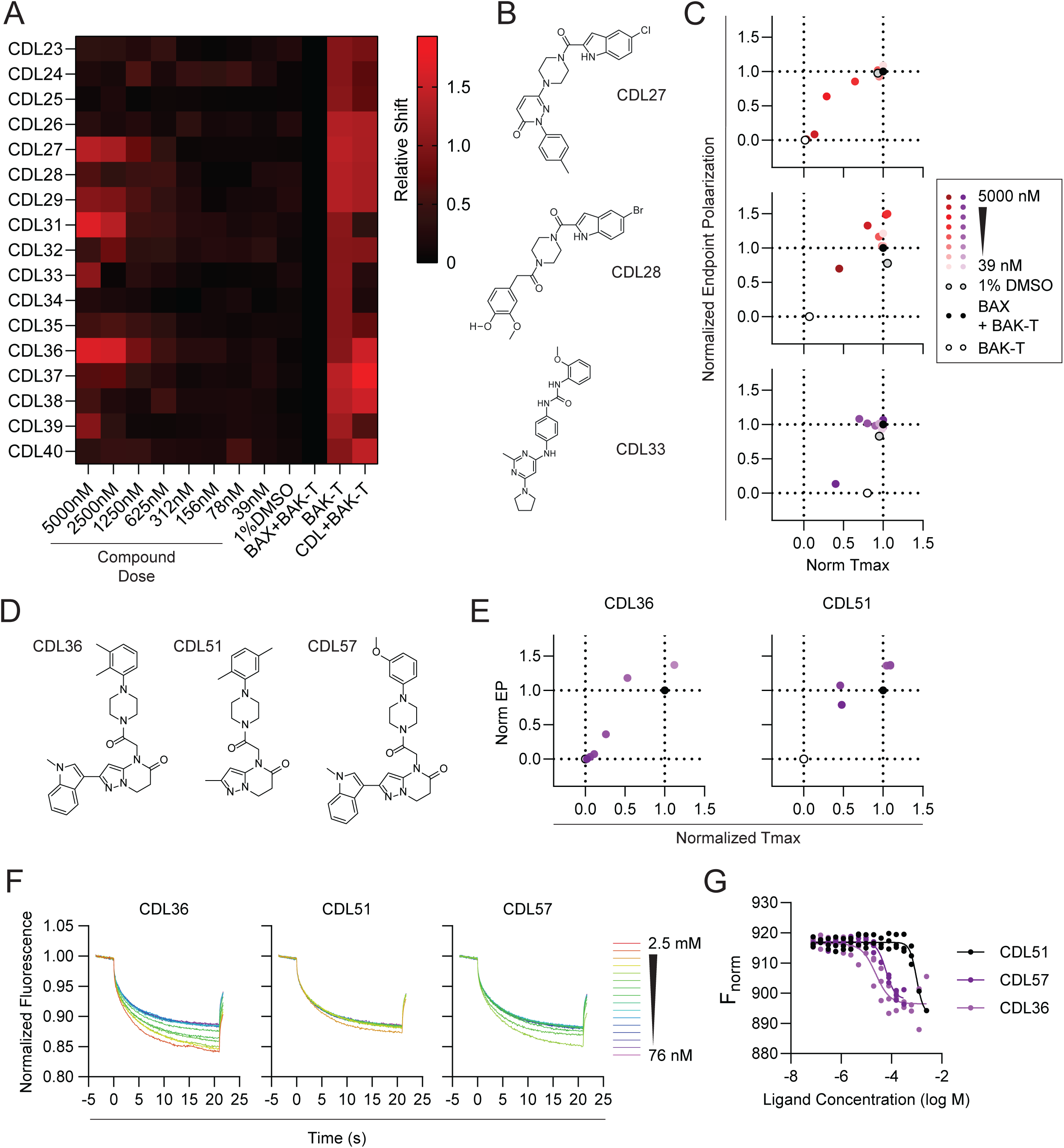
Candidate inhibitors bind directly to recombinant pore-forming proteins. A. Heatmap depicting relative shift in FLAMBE parameters for 19 candidate inhibitor compounds, indicative of conformational change induced by compound binding to BAX. Relative shift is calculated based on the distance (as depicted in C) from the point representing untreated polarization of TAMRA-labeled BAK BH3 peptide along two axes: EP (endpoint fluorescence polarization) and Tmax (time of maximum fluorescence polarization). Recombinant BAX concentration is 90 nM. Data represent average of two technical replicates. B. Structures of small molecules characterized in C and in Figure 1F-G. C. Plots depicting EP and Tmax across doses of the compounds in B, relative to BAK-TAMRA plus BAX and BAK-TAMRA alone. Recombinant BAX concentration is 90 nM. Data represent average of two technical replicates. D. Structures of small molecule CDL36 and two analogs identified from the primary screen as structurally similar. E. Plots depicting EP and Tmax across doses of the compounds in D. Recombinant BAX concentration is 65 nM. Data represent average of two technical replicates. F. Microscale thermophoresis assay showing normalized fluorescence of fluorescent-labeled recombinant BAX in the presence of indicated doses of the candidate compounds or controls over time. Infrared laser was applied from t = 0 to 21 seconds. G. Plot of normalized fluorescence, expressed as 1000X the ratio between the heated and cold states, across CDL36 concentrations.

To better understand the chemical properties of CDL36, we identified two structurally similar compounds, CDL51 and CDL57, from the primary screening library (Figure 2D) and obtained fresh stocks for comparison in activity and binding assays. To test their activity, we quantified the induction of MOMP by BIM peptide in the presence or absence of the candidate inhibitors as measured by loss of cytochrome *c* staining. CDL51, which lacks the bicyclic aromatic indole group found in CDL36, showed no ability to prevent peptide-induced MOMP (Figure S2C). By contrast, CDL57, which differs from CDL36 in the substituents on its benzene ring, showed only a modest reduction in activity. FLAMBE analysis of CDL36 and CDL51 demonstrated patterns of BAX binding consistent with the measured activities of the compounds; application of CDL36 in the recombinant BAX system yielded a strong, dose-dependent decrease in fluorescence polarization, while CDL51 did not (Figure 2E).

As an orthogonal approach to the assessment of BAX binding, we also employed microscale thermophosesis (MST) to measure BAX binding for our most potent candidate inhibitors and control compounds. The MST assay measures changes in motility of a fluorescently labeled recombinant target protein in solution upon binding of an inhibitor. We produced recombinant BAX (rBAX; Figure S2D-E)^23^ and confirmed the ability of the recombinant protein to restore BIM peptide sensitivity to permeabilized BAX/BAK double knockout HeLa cells, which are otherwise insensitive due to the absence of these pore-forming proteins. At 1 nM and 500 pM doses, BAX activity (as measured by cytochrome *c* release) was dependent on BIM peptide co-treatment; at higher doses, BAX alone was sufficient to induce MOMP (Figure S2F). Having confirmed that rBAX behaved as expected in cellular assays, we labeled the protein for fluorescence detection in MST assays. MST analysis confirmed the binding of CDL36 to rBAX with an EC_50_ of 23.6 µM (Figure 2F-G, 95% CI 14.7-37.8 µM). The affinity of CDL57 for rBAX was moderately weaker than CDL36, and CDL51 showed the weakest binding of the three. Thus, *in vitro* binding assays suggested that CDL36 binds directly to BAX in solution, and the strength of this interaction can be modulated via modifications to the small molecule structure of CDL36.

### CDL36 Prevents Induction of MOMP by Both BAX and BAK

To test the necessity of BAX and BAK for the protective effect of CDL36 across distinct cell line backgrounds, we generated BAX and BAK single and double CRISPR knockout lines from A549 lung adenocarcinoma and HT-1080 fibrosarcoma cells (Figure 3A, S3A-C). Preceding experiments had primarily tested the activity of CDL36 in permeabilized cells and *in vitro* systems over short (up to 1 h) timescales; therefore, before performing long-term cytotoxicity assays, we first conducted assays in intact cells where pro-apoptotic signaling was induced over similar timescales. First, we tested the induction of MOMP by the combined inhibition of BCL-xL and MCL-1 in unpermeabilized cells, measuring the ability of CDL36 and BAI1 to block BH3 mimetic-induced MOMP. We measured MOMP by staining with tetramethylrhodamine ester (TMRE), a dye whose fluorescence is lost upon MOMP-associated mitochondrial depolarization^24^. After 1 hour of treatment with the combined BH3 mimetics A-1331852 (a BCL-X_L_ inhibitor) and S63845 (an MCL-1 inhibitor) in the presence or absence of the inhibitors, we measured MOMP via TMRE staining and FACS analysis (Figure 3B, S3D). We observed that BH3 mimetic treatment caused a dose-dependent loss of mitochondrial membrane potential in CRISPR control cells and some BAX or BAK knockout cells in both the A549 and HT-1080 background, but not BAX/BAK double knockouts of either cell type. The loss of membrane potential was prevented by both CDL36 and BAI1 in all sensitive cell lines, indicating that neither BAX nor BAK was individually required for protection against MOMP by CDL36 or BAI1 and that both compounds were protective against MOMP induction in intact cells in culture.

**Figure 3.**
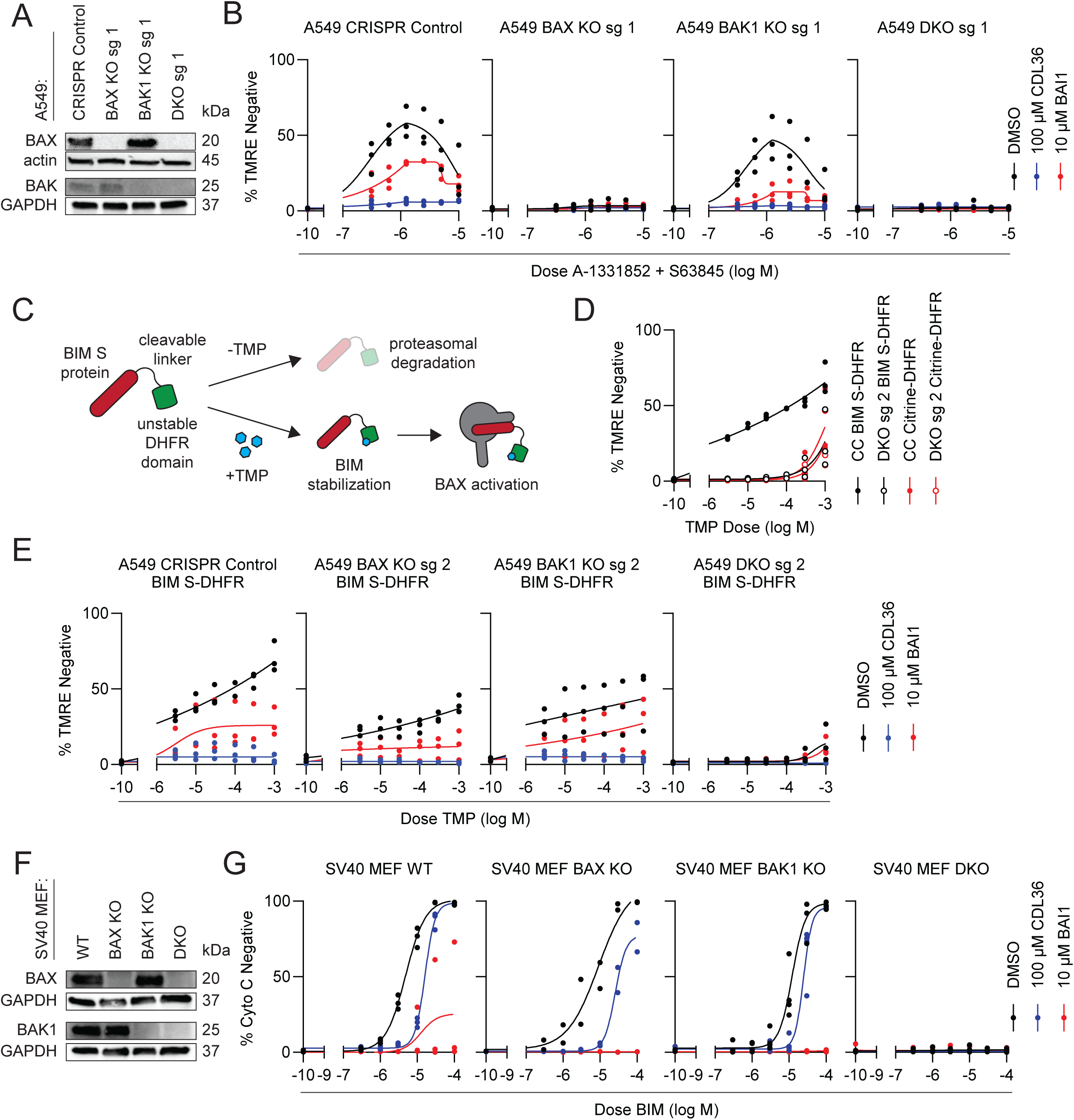
CDL36 and BAI protect against MOMP in cells with intact plasma membranes. A. Western blot analysis of knockout and control A549 lung adenocarcinoma cells generated by CRISPR-mediated disruption of the *BAX* and *BAK1* loci. B. CDL36 and BAI1 activity as measured by tetramethylrhodamine ester (TMRE) signal loss as an indicator of MOMP. Cells were co-treated for 1 h with a range of doses of the combined BCL-xL inhibitor A-133852 and MCL-1 inhibitor S63845 and the indicated inhibitor dose or vehicle control. A biphasic sigmoidal curve was fit to the data to account for the peak in TMRE at intermediate inhibitor doses. n = 3 biological replicates. C. Schematic depicting the mechanism for induction of BIM protein and subsequent MOMP using the dihydrofolate reductase (DHFR) fusion system. D. BAX/BAK- and BIM-dependent induction of MOMP by a range of trimethoprim (TMP) doses in cells expressing the DHFR fusion system. The DHFR domain was fused either to BIM or Citrine as a control, and the fusion construct was expressed in either BAX/BAK DKO or CRISPR control cells. Cells were treated with TMP for 1 h. n = 3 biological replicates. E. CDL36 and BAI1 activity as measured by TMRE stain for MOMP upon BIM induction. BIM expression was induced by 1 h treatment with a range of TMP doses, co-treated with the indicated inhibitor dose or vehicle control. n = 3 biological replicates. F. Western blot analysis of wild-type and *Bax* and/or *Bak1* knockout SV40 mouse embryonic fibroblasts (MEFs). G. CDL36 and BAI1 activity in wild-type and knockout MEFs as measured by BH3 profiling. Cytochrome c staining is an indicator for MOMP upon treatment with a range of BIM peptide doses, co-treated with the indicated inhibitor dose or vehicle control. n = 3 biological replicates.

To account for potential off-target effects of BH3 mimetics, we also implemented an orthogonal method for the rapid induction of MOMP that is not reliant on direct small molecule interactions with BCL-2 family proteins (Figure 3C). In this system, mouse BIM protein fused to an engineered *E. coli* dihydrofolate reductase (DHFR) domain is constitutively expressed in lentivirally transduced cells. The DHFR domain is unstable, leading to proteasomal degradation of the BIM-DHFR fusion^25^. Treatment with trimethoprim (TMP), a small molecule inhibitor of DHFR, stabilizes the fusion construct, rapidly upregulating BIM-DHFR levels in a dose-dependent fashion. Although BIM-DHFR fusion is leaky, yielding heightened levels of basal BIM expression (Figure S3E), we validated that MOMP induction in BIM-DHFR expressing cells (as measured by TMRE signal loss) is dependent on TMP treatment and the presence of BAX or BAK (Figure 3D, S3G). TMP treatment alone does not induce MOMP, as demonstrated by the retention of TMRE signal in TMP-treated cells expressing a Citrine-DHFR fusion in place of BIM-DHFR. The protective effects of CDL36 and BAI1 against TMP-induced MOMP in CRISPR control and BAX/BAK single and double knockouts was consistent with the effects observed in BH3 mimetic-induced MOMP (Figure 3E), indicating that the compounds’ effects are not restricted to MOMP induced by direct inhibitors of anti-apoptotic BCL-2 family proteins. The structural analogs CDL51 and CDL57 showed activity in the TMP-based assay consistent with their activity against mimetic induced MOMP (Figure S3H), further validating the relationship between structure, BAX binding, and MOMP-inhibitory activity.

Lastly, to test the species specificity of CDL36, we utilized BAX and BAK knockout mouse embryonic fibroblasts (MEFs) to test the activity of CDL36 in mouse cells in culture (Figure 3F). In BH3 profiling experiments measuring cytochrome *c* loss upon pro-apoptotic peptide treatment, we observed that both CDL36 and BAI1 reduced the induction of MOMP in wild type and both BAX and BAK KO MEFs (Figure 3G). Consistent with the human cell line data, neither BAX nor BAX was individually required for the protective effect of either CDL36 or BAI1 in MEFs, suggesting that these compounds might inhibit pore formation by either protein in mouse cells as well. The relative activity of CDL36 and BAI1 differed in MEFs relative to the tested human cell lines, suggesting a higher relative potency of BAI1 in mouse vs. human cells. On balance, however, our results indicate that CDL36 has broad MOMP-inhibitory activity across human and mouse cell types and against distinct pro-apoptotic stimuli.

### Activity of CDL36 is Specific to BAX/BAK-Dependent Death and Does Not Require VDAC2

Because we had measured the activity of CDL36 and BAI1 using TMRE, a stain for mitochondrial membrane potential, we next sought to confirm that the observed effects were due to a blunting of cell death and not a non-specific effect on mitochondrial membrane integrity or bioenergetics. We utilized the apoptosis inducer raptinal, which has been reported to induce apoptosis through direct activation of caspases or other mechanisms not requiring BAX/BAK pore formation^26^. Consistent with these reports, we found that raptinal treatment induced TMRE loss in a dose dependent manner in both CRISPR control and BAX/BAK single or double knockout A549 cells (Figure 4A). Raptinal treatment also induced phosphatidylserine (PS) externalization, a later event in the apoptotic cascade, as measured by staining with Annexin V (Figure S4A), but neither TMRE loss nor Annexin V positivity was abolished by cotreatment with CDL36 or BAI1. Annexin V staining was limited after 1 hour of treatment, so we tested an additional timepoint at 24 hours after raptinal application, reducing the dose of CDL36 to 25 µM to avoid raptinal-independent toxicity observed at timepoints beyond 1 hour. We observed strong raptinal-induced Annexin V (Figure 4B) and TMRE (Figure S4B) signals across all genotypes after 24 hours, but no inhibition of death signaling by CDL36 or BAI1.

**Figure 4.**
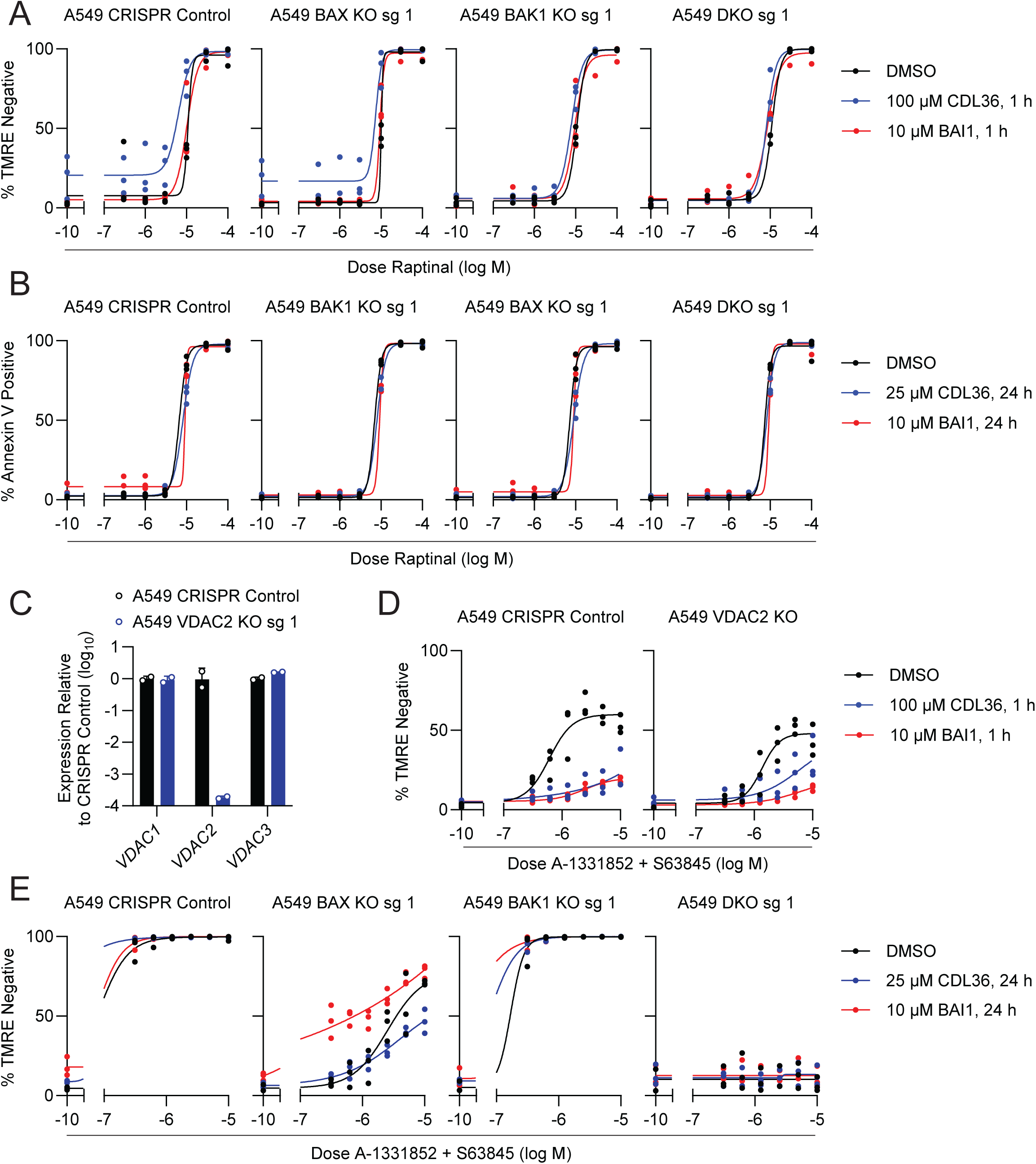
Protective effect of CDL36 against MOMP is specific but transient. A. CDL36 and BAI1 activity as measured by TMRE signal loss as an indicator of MOMP. Cells were co-treated for 1 h with a range of doses of the BAX/BAK-independent apoptosis inducer raptinal and the indicated inhibitor dose or vehicle control. n = 3 biological replicates B. CDL36 and BAI1 activity as measured by Annexin V staining for PS externalization as an early indicator of apoptotic cell death. Cells were co-treated for 24 h with a range of doses of raptinal and the indicated inhibitor dose or vehicle control. n = 3 biological replicates. C. Confirmation of CRISPR-mediated knockout of VDAC2 by qPCR in A549 cells. n = 2 technical replicates. D. CDL36 and BAI1 activity in CRISPR control and VDAC2 KO A549 cells and WT and VDAC2 KO HCT116 cells as measured by TMRE. Cells were co-treated for 1 h with a range of doses of combined A-133852 and S63845 and the indicated inhibitor dose or vehicle control. n = 3 biological replicates. E. CDL36 and BAI1 activity as measured by TMRE signal loss at 24 h after treatment. Cells were co-treated with a range of doses of combined A-133852 and S63845 and the indicated inhibitor dose or vehicle control. N = 3 biological replicates.

Some recently developed small molecule modulators of BAX/BAK protein activity have been reported to act through disruption of the interaction between BAX or BAK and VDAC2, a mitochondrial outer membrane protein whose interactions with BAX and BAK can modulate the activation of these proteins^27^. We generated knockouts of VDAC2 in an A549 background to test the protective activity of CDL36 and BAI1 in cells lacking VDAC2. Because antibodies against VDAC2 can also recognize the other VDAC isoforms, we validated these knockouts via qPCR, demonstrating that the VDAC2 CRISPR knockout specifically disrupted VDAC2 expression, but not VDAC1 or 3 expression, at the mRNA level (Figure 4C). To test the activity of CDL36 and BAI1 in this context, we measured MOMP via TMRE staining upon BH3 mimetic treatment (Figure 4D). Although we did not carry out the necessary control experiments to conclusively measure the effect of VDAC2 loss on apoptotic sensitivity overall, we noted that the selected VDAC2 knockout clone showed decreased sensitivity to BH3 mimetics relative to CRISPR control cells. When combining CDL36 or BAI1 with the BH3 mimetic treatment, we observed that both compounds effectively prevented BH3 mimetic-induced TMRE loss in both the VDAC2 knockout and CRISPR control cells. Our results indicated that CDL36 specifically protects against BAX/BAK-dependent death in a manner that does not require VDAC2; we next sought to assess the durability of this protection in longer term assays of cell death.

### Protection Against BH3 Mimetic-Induced MOMP by CDL36 and BAI1 is Transient

To better understand the kinetics of the MOMP-inhibitory effects of CDL36, we tested its ability to prevent BH3 mimetic-induced MOMP after 4 and 24 hours of treatment in A549 cells, comparing BAX, BAK, and double knockout cells to CRISPR controls. At 4 hours, CDL36 and BAI1 showed only modest protection against BH3 mimetic-induced MOMP across a broad range of doses compared to that observed at 1 hour (Figure 4E, S4C-E). As observed in the 4 hour raptinal treatment, 100 µM CDL36 showed signs of BH3 mimetic-independent toxicity at 4 hours; we therefore tested CDL36 at 25 µM at 24 hours. At 24 hours, neither CDL36 nor BAI1 substantially protected mimetic-treated cells against MOMP or PS externalization, supporting the conclusion that these compounds’ protective effects are transient in A549 cells treated with BH3 mimetics. We therefore set out to test whether death induced by other perturbations might be more durably inhibited by CDL36, or if protection was similarly transient across lethal stimuli.

### CDL36 Specifically Protects Against Doxorubicin-Induced Death

To test the ability of CDL36 to promote longer-term protection against various lethal stimuli, we performed live cell time lapse imaging of cell death in CRISPR control and BAX knockout A549 cells treated with a panel of cytotoxic compounds. We first tested a range of doses of various compounds to identify doses high enough to induce death even in BAX KO cells (data not shown), reasoning that some protective activity might be gained or lost in this genotype relative to the control cells. We selected doses at which to test 8 lethal stimuli, combining each with CDL36 or a DMSO vehicle control for a 72 hour treatment. At each timepoint, we calculated the Lethal Fraction, a metric quantifying dead cells as a fraction of total cell number^28^, based on SYTOX green- and Hoechst-positive object counts. For the full timecourse, we then calculated the area under the curve (AUC) for the Lethal Fraction metric over time (Figure 5A). In conditions without other lethal stimuli, 50 µM CDL36 induced substantial death in both WT and BAK KO cells (Figure 5B, S5A); we therefore limited our subsequent analyses to the 25 and 12.5 µM doses of CDL36.

**Figure 5.**
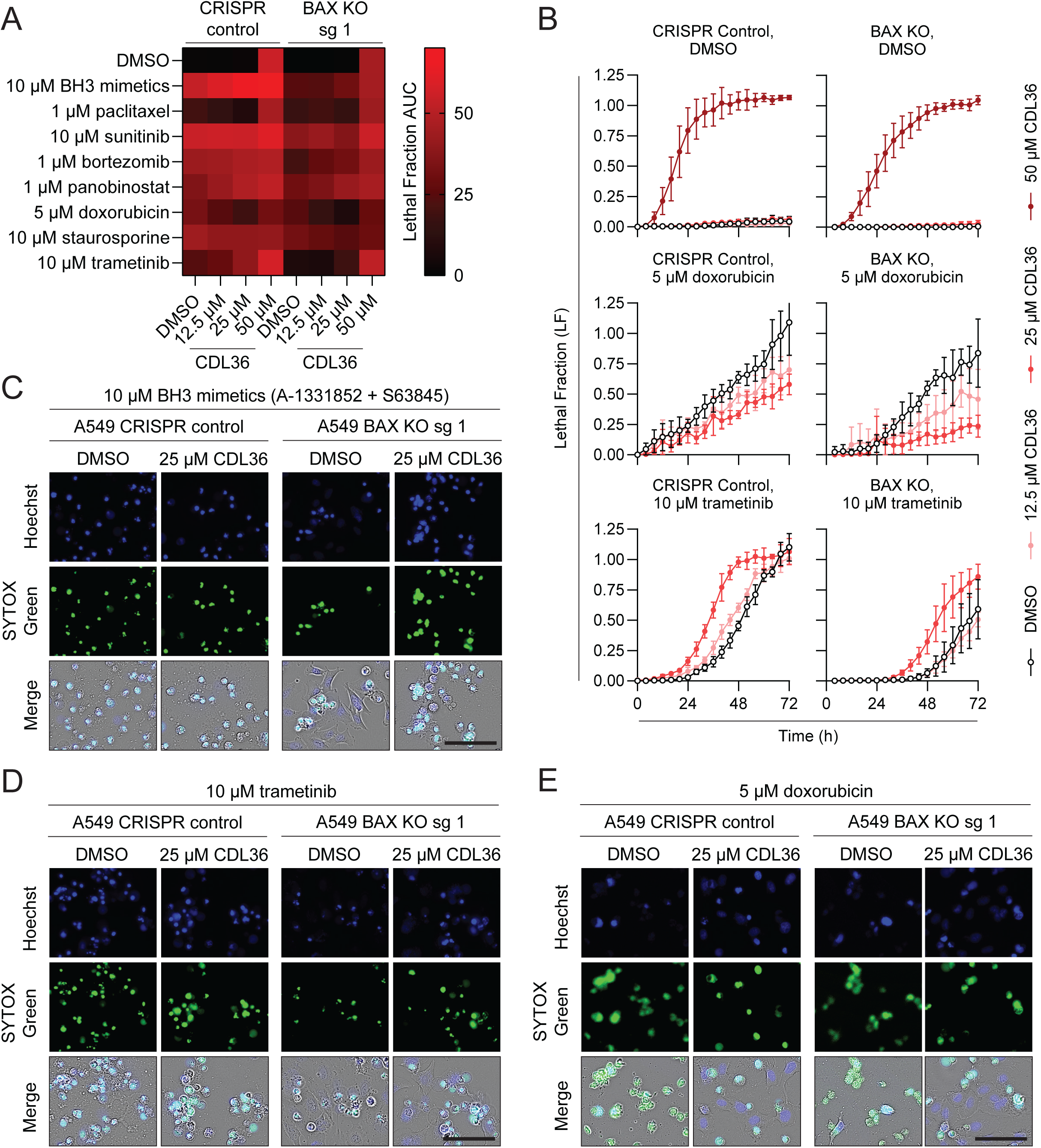
Among a panel of cytotoxic compounds, doxorubicin-induced death is specifically inhibited by CDL36. A. Heatmap depicting the area under the curve (AUC) calculated from the Lethal Fraction (LF) metric of cell death over 72 hours of live cell timelapse imaging. CRISPR control or BAX KO A549 cells were treated with the indicated doses of 8 lethal stimuli or vehicle control, combined with CDL36 doses from 12.5 to 50 µM. Data represent the mean of 3-4 biological replicates. B. Example LF timecourses comparing death over time in response to DMSO, doxorubicin, or trametinib in the presence or absence of the indicated CDL36 doses. Data represent the mean and SD of 3-4 biological replicates. C-E. Representative images depicting total (Hoechst+) or dead (SYTOX Green+) cells after 48 hours of treatment with 10 µM combined BH3 mimetics (C), 10 µM trametinib (D), or 5 µM doxorubicin (E) in the presence or absence of 25 µM CDL36. Scale bar = 100 µm.

For most lethal stimuli, addition of CDL36 had little effect on the observed death or, as in the case of trametinib, increased the rate or degree of death over time (Figure 5B-D, S5B). By contrast, death induced by doxorubicin was substantially reduced when combined with CDL36 in either WT or BAX KO cells. Cells treated with the combination of doxorubicin and CDL36 underwent proliferative arrest and displayed the associated changes in morphology (Figure 5E, S5C) but the combined treatment yielded substantially less cell death than doxorubicin treatment alone over the tested timecourse. Thus, for specific lethal stimuli, CDL36 is capable of preventing or delaying cell death over multi-day timescales.

## Discussion

Our results demonstrate the utility of the BH3 profiling methodology in a high-throughput screening application for the first time, allowing us to identify a series of small molecules with cytoprotective activity in various contexts. In characterizing these screening hits, we identified a subset showing direct binding to BAX, including our most potent candidate inhibitor, CDL36. The transient anti-apoptotic activity of CDL36 illustrates the outstanding challenges in developing and applying inhibitors of BAX, BAK, and other BCL-2 family proteins. Previous efforts to inhibit BAX and BAK have highlighted the importance of stoichiometry between inhibitors and targets, with the same molecules showing BAX/BAK activation at low doses, but inhibition at high doses^16^. This is consistent with the mechanism by which BIM and other activator proteins activate the pore-forming BCL-2 family proteins, acting by a “hit-and-run” mechanism that does not require sustained binding of activators to BAX or BAK^29–31^ and may therefore require inhibitors to be present in excess and continuously to prevent BAX/BAK activation.

Other investigators have highlighted the complex factors driving stabilization or destabilization of pore-forming BCL-2 family proteins in diverse conformations and complexes, whereby peptide binders of BAK may promote either its inhibition or activation depending on factors beyond simple binding affinity^15^. Although the present implementation is limited by leaky expression, we anticipate that the BIM-DHFR fusion system presented here might prove useful in untangling these factors, allowing iterative characterization of key residues for protein-protein interactions in intact cells. As protein degron systems undergo continued development and refinement^32,33^, the combined insights from studies employing small molecule and peptide binders will enable a more comprehensive mechanistic characterization of BAX/BAK activation.

Small molecule modulation of BAX and BAK activation is further complicated by the structural similarity between BCL-2 family proteins and the resulting potential for pleiotropy in binding by inhibitors of this family^34,35^. Putative BAX/BAK inhibitors, including CDL36, might bind to anti-apoptotic BCL-2 family proteins at some doses and timepoints, potentially contributing to the cytotoxic or mixed effects observed here. More extensive MST, LUV studies, and other binding assays will be required to determine the full complement of small molecule binding affinities to the full set of BCL-2 family proteins. With a sufficiently specific binder of BAX or BAK, modulation of apoptosis might be an ideal application for proteolysis-targeting chimeras (PROTACs) that could promote ubiquitin proteasome-mediated degradation of BAX or BAK even without complete inhibition of the pool of pore-forming BCL-2 family proteins within the cell^36^. This approach has previously been successfully applied to the targeting of BCL-2 family proteins to induce apoptosis in cancer models^37,38^ and has recently even advanced to the clinic^39^.

Whether through inhibition or degradation, the modulation of BAX and BAK activity has important potential applications for both laboratory investigation and clinical abrogation of cell death. Doxorubicin-induced cardiotoxicity is a well-characterized and dose-limiting side effect in cancer patients receiving this therapy^40^, especially in children^41^, and elucidating the mechanism of this toxicity may enable the development of therapies with improved efficacy against tumor cells or reduced toxicity in healthy cardiomyocytes and other cells. The roles of BAX and BAK in normal and cancerous cells require continued investigation, and novel and newly characterized chemical tools will enable this investigation. Notably, the protection by CDL36 against doxorubicin-induced death in the present study was most durable in cells lacking BAX; if this finding holds across broader cellular and tissue contexts, it might suggest that protection by this mechanism could be particularly potent in cardiomyocytes, neural cells, and other cell types where the expression of BAX and BAK in adult mammals is quite low^42–45^.

In general, the chemical modulation of BAX and BAK remains an important goal for investigators of cell death. Some applications, such as in radiation-induced cell death, are well-validated to involve BAX/BAK-mediated apoptotic cell death^46^ and protection from apoptosis by blocking these effectors has been shown to protect against short- and long-term tissue dysfunction driven by radiation exposures^43,47^. For other potential applications, including in neurodegeneration, defining the contribution of apoptosis or other BAX/BAK-dependent pathways has been difficult in the absence of robust small molecule BAX/BAK inhibitors^48^. The eventual clinical application of apoptosis inhibitors will require careful consideration of expected side effects, including the potential for promotion of tumorigenesis and autoimmunity with sustained and systemic inhibition^1^. With these considerations in mind, however, the further development of potent and specific cell death inhibitors will expand the small molecule toolbox for the continued study of BAX, BAK, and other critical mediators of cell death.

## Acknowledgments

Small molecule screening and data deposition were performed with the support of the ICCB-Longwood Screening Facility at Harvard Medical School. Microscale thermophoresis was performed with the support of the Center for Macromolecular Interactions at Harvard Medical School. FACS was performed with support of the Dana-Farber Cancer Institute Flow Cytometry Core Facility. We thank Maryam Ibrahim for assistance with experiments and Dr. Grant Dewson at the Walter and Eliza Hall Institute for generously sharing SV40 MEF cells with our laboratory.

## Funding

This work was supported by funding from:

National Institutes of Health grant R00CA188679 (KAS)

Blavatnik Biomedical Accelerator at Harvard University (KAS)

National Institutes of Health grant K99AG086405 (ZI)

## Resource Availability

### Lead Contact

Requests for information and materials should be directed to the corresponding author, Kristopher A. Sarosiek, and will be fulfilled whenever possible.

### Data Availability

Primary screening data will be made available via PubChem (pubchem.ncbi.nlm.nih.gov) as a BioAssay. All other data generated in this study are available upon request from the corresponding author.

### Materials Availability

Plasmids produced for the work described will be made available via Addgene (addgene.org). Where possible, cell lines will be made available by the authors upon request.

## Supplementary Materials

Supplementary Figures:

Figure S1 (Related to Figure 1)

Figure S2 (Related to Figure 2)

Figure S3 (Related to Figure 3)

Figure S4 (Related to Figure 3)

Figure S5 (Related to Figure 3)

Supplementary Tables:

Table S1: Description of Small Molecule Libraries

Table S2: Oligonucleotide Sequences

Table S3: PCR and qPCR Primer Sequences

## Materials and Methods

### Cell Lines and Culture Conditions

Cell lines were cultured in high glucose DMEM base media (Thermo Fisher Scientific, cat #11965118) supplemented with 10% fetal bovine serum (Thermo Fisher Scientific, cat #A5256701) and 1% penicillin/streptomycin (Fisher Scientific, cat #15-140-163). Cell line identity was verified by STR profiling (Lab Corp, Burlington, NC) and cells were tested regularly for mycoplasma using a MycoAlert Plus luminescent assay (Lonza, cat #LT07-710). Cells were incubated at 37°C at 5% CO_2_, passaging as needed using Trypsin-EDTA (Thermo Fisher Scientific, cat #25200114). Cell counting for passaging and seeding was performed with a Beckman Coulter Vi Cell XR.

### Generation of CRISPR-Mediated Knockout Cell Lines

Single guide RNA (sgRNA) expression plasmids were generated from the pSpCas9(BB)-2A-GFP vector (Addgene plasmid #48138, a gift from Feng Zhang) following the published protocol^49^. Briefly, two sgRNAs per target gene were selected using the Synthego guide design tool, and complementary oligonucleotides were designed containing the guide sequences and compatible overhangs (Table S2). Primer pairs for genomic DNA amplification were designed using the NCBI Primer BLAST tool to encompass the cut sites for each target gene (Table S3). 1 µg of the vector plasmid DNA was digested in a 20 µL reaction using Fast Digest BpiI and FastAP Thermosensitive Alkaline Phosphatase (Thermo Fisher Scientific), and the product was gel purified using a Monarch DNA Gel Extraction Kit (New England Biolabs). Each oligonucleotide pair was phosphorylated and annealed in a 10 µL reaction containing 10 µM of each oligonucleotide, 1X T4 DNA Ligase Reaction Buffer (New England Biolabs), 0.5 µL T4 Polynucleotide Kinase (New England Biolabs), and nuclease-free water to volume. Reactions were incubated at 37°C for 30 min, followed by 95°C for 5 min, and cooled to 25°C at a rate of 5°C/5 min in a thermal cycler. Annealed oligonucleotides were diluted 1:200 and ligated into the digested pSpCas9(BB)-2A-GFP backbone in a 10 µL reaction containing 50 ng digested plasmid DNA in a Quick Ligation Reaction (New England Biolabs, cat #).

For transfection, cells were seeded at a density of 3*10^5^ cells per well in 6-well plates and cultured for 24 h prior to transfection using 7.5 µL Lipofectamine 3000 (Thermo Fisher Scientific) and 2.5 µg plasmid DNA per transfection. On the day of transfection, 96-well plates for single-cell sorting were prepared containing DMEM media supplemented with 20% fetal bovine serum and 1% penicillin/streptomycin and stored in a tissue culture incubator until sorting. 24 hours post-transfection, cells were dissociated with trypsin, passed through a cell strainer to obtain a single-cell suspension, and sorted by FACS into the prepared plates at a density of one cell per well at the Dana Farber Cancer Institute Flow Cytometry Core. GFP-positive cells were identified, setting the gate to exclude autofluorescent events. Following clonal expansion, genomic DNA was extracted from candidate clones using QuickExtract DNA Extraction Solution (VWR, cat #76081-768), amplified using the designed PCR primers, and sequenced to identify clones containing insertions or deletions at the intended cut site (Plasmidsaurus, Boston, MA). Candidate clones were further characterized by Western blotting and/or qPCR as described below.

### Generation of DHFR Fusion Constructs

Plasmids PB-PGK-Cit-tevs-DHFR (plasmid #116040, a gift from Michael Elowitz), pCMV-Tag2B Flag BimEL (plasmid #23090, a gift from Roger Davis), and pLX_TCR311 (plasmid #113668, a gift from John Doench) were obtained from Addgene. To facilitate subsequent cloning reactions, the Cit-tevs-DHFR sequence was cloned into a Gateway entry vector. PCR primers were designed to amplify the Cit-tevs-DHFR sequence with flanking attB sites (Table S3). PCR amplification was performed using Q5 Hot Start High-Fidelity DNA Polymerase (New England Biolabs, cat #M0494S) and the attB PCR product was cloned into the pDONR221 plasmid (Thermo Fisher Scientific, cat # 2536017) in a BP clonase reaction according to manufacturer instructions (Thermo Fisher Scientific, cat #11789020). The resulting pENTR_Cit-tevs-DHFR plasmid was used for insertion of the Bim EL coding sequence in place of the Citrine sequence via restriction cloning. Primers were designed to amplify the BimEL coding sequence from pCMV-Tag2B Flag BimEL, appending a 3’ BamHI sequence and 5’ XhoI sequence (Table S3). pENTR_Cit-tevs-DHFR backbone and BimEL PCR product were digested with BamHI and XhoI (New England Biolabs, cat #R3136S, R0146S), gel purified, and ligated in a Quick Ligation reaction. The resulting plasmid (pENTR_BimEL-tevs-DHFR), was then used to generate a lentiviral expression plasmid with the pLX_TRC311 backbone via LR clonase reaction (Thermo Fisher Scientific, cat #11791020). The lentiviral expression plasmid (pLX_TRC311_BimEL-tevs-DHFR) and control Citrine plasmid (pLX_TRC311_Cit-tevs-DHFR) were used to generate lentivirus for transduction as described below.

### Generation of Lentivirus and Infection/Selection of Cell Lines

To generate lentivirus, HEK293T cells were seeded at a density of 2.5*10^5^ cells per well in a 6 well plate and allowed to adhere and recover for 24 hours before transfection. Transfections were carried out using 7.5 µL Lipofectamine 3000 (Thermo Fisher Scientific), the lentiviral expression vector, and the requisite 3rd generation packaging plasmids, pCMV-VSV-G, pMDLg/pRRE, and pRSV-Rev (Addgene plasmid #8454, a gift from Bob Weinberg, and #12251 and #12253, gifts from Didier Trono). Virus-containing media was harvested once after 48 hours, the media was refreshed, and a second harvest was performed after an additional 24 hours. The combined virus-containing media was passed through a .22 µm PVDF filter, aliquoted, and stored at -80°C. For infection, host cell lines were seeded in 24-well plates at a density of 50k per well and allowed to adhere for 24 hours. Infection media was prepared combining virus media with polybrene, applied to the cells, and incubated for 24 hours, after which the infection media was removed and replaced with standard culture media for 24 hours. The infected cells were expanded and selected with an appropriate antibiotic.

### Primary Small Molecule Screen

Chemical libraries were obtained by the ICCB-Longwood core screening facility; a total of 44,827 compounds were screened from various libraries (Table S1). The screen was run in two technical replicates of each of two lymphoma cell lines, U937 and OCI-Ly1. A separate screen, comprising *BAX*/*BAK1* double knockout (DKO) HeLa cervical cancer cell lines, was used to identify potential autofluorescent compounds. On days screening was performed, U937 or OCI-Ly1 cells were suspended in Mannitol Experimental Buffer (MEB; 150mM Mannitol, 10mM HEPES, 50mM KCl, 20µM EGTA, 20µM EDTA, 0.1% BSA, 10mM Succinate, pH 7.5) at a density of 4 million cells/mL. The cell suspension was mixed with an equal volume of MEB buffer containing 0.015% w/v digitonin and 30 µg/mL oligomycin to permeabilize cells. For treatment with library compounds, 30 µL of the permeabilized cell mixture was added to each well of a 384 well plate (Greiner, cat #781076) using a Multidrop Combi Reagent Dispenser (Thermo Fisher Scientific), resulting in a final cell number of 60,000 cells/well. The library compounds (most at 2 mg/mL, 5 mg/mL, or 10 mM doses) were added to plates at a volume of 33 nL by stainless steel pin array using a custom robotic transfer platform (ICCB-Longwood), and plates were incubated for 25 minutes at 30 degrees C.

For peptide treatment and staining, a mixture was prepared in MEB buffer containing 3 mM JC-1, 15 mM β-mercaptoethanol, and 3 uM each BIM and BID peptide. The treatment/staining mixture was added to cells at a volume of 15 µL/well using the Multidrop Combi, resulting in a final concentration of 1 µM for each peptide. Cells were incubated for 60 minutes at 30 degrees C before measurement of mitochondrial outer membrane permeabilization (MOMP) via JC-1 fluorescence. Membrane potential-dependent red fluorescence of JC-1 J-aggregates was measured via excitation at 545 nm and emission at 595 nm using a Synergy H1 Multi-Mode Plate Reader (Biotek). Z-scores were calculated on a per-plate basis based on the mean and standard deviation of fluorescence readings for all experimental wells, and z-scores were used to select compounds for follow-up secondary screening as described below.

### Secondary Screening

Hits from the primary screen were identified as compounds with a z-score greater than 3 for fluorescence values for both replicates in either U937, OCI-Ly1, or both without an effect on HeLa DKO cells (z-score less than 1). Libraries of compounds meeting these criteria were selected for secondary screening in three groups: compounds scoring as hits in both lymphoma lines (116 screening wells, 115 compounds – 2 wells contained different doses of the same compound), compounds scoring as hits in U937 only (89 compounds), or compounds scoring as hits in OCI-Ly1 only (100 compounds). Rescreening was conducted by cotreating U937 cells (dual or U937 hits) or OCI-Ly1 cells (OCI-Ly1 hits) with the library compounds at the original screening dose and BIM peptide at 0.3 and 0.03 µM in 96 well plates in three biological replicates. Peptide treatment, fixation, quenching, and staining were performed according to the protocol for BH3 profiling described below. Compounds which yielded the lowest percentages of cytochrome *c* negative cells, blocking the pro-MOMP activity of BIM, were selected for further testing against a full dose response of BIM peptide treatment.

For the full BIM dose response, candidate compounds were ordered from ChemDiv, resuspended in DMSO to generate 50 mM stock solutions, and again tested in three groups: dual (11 compounds), U937 (8 compounds), and LY1 (2 compounds) hits. Compounds were tested at 100, 10, and 1 µM in combination with BIM concentrations from 0.1 to 100 µM in a 7 point, half-log dose response, or with a DMSO vehicle control.

### Western Blotting

Cells were harvested from culture, centrifuged at 500 x g for 5 minutes, and washed via resuspension in 750 µL of PBS (137 mM NaCl, 10 mM phosphate, 2.7 mM KCl, pH 7.4). The cells were lysed with a RIPA lysis buffer, (50 mM Tris-HCl, 150 mM NaCl, 1% NP-40, 0.05% sodium deoxycholate, 0.02% SDS; Boston BioProducts, cat #BP-115-5X) supplemented with cOmplete Protease Inhibitor (Millipore Sigma, cat #4693132001) and Phosphatase Inhibitor (Thermo Fisher Scientific, cat # A32957), for 1 hour on ice. The lysate solution was centrifuged (10,000 x g, 10 minutes) before being transferred to clean microcentrifuge tubes and stored at -20 °C. Protein content of the lysates was quantified via BCA Assay (Thermo Fisher Scientific, cat #23209). Samples for SDS-PAGE were prepared with NuPAGE LDS Sample Buffer (Thermo Fisher Scientific, cat #NP0007) at a protein concentration of 1 mg/mL protein, supplemented with 50 mM DTT (GoldBio, cat #DTT10). In a 10 well any-kD SDS-PAGE gel (Bio Rad, cat # 4569033), 20 µg of protein was loaded and run at 120 V for 1 hour, then transferred to a PVDF membrane (BioRad, cat #1620177). The membranes were incubated for 30 minutes in blocking buffer (10.1 mM Na2HPO4, 1.76 mM KH2PO4, 2.7 mM KCl, 137 mM, 0.1% Tween 20, 5% skim milk powder). Primary antibodies were diluted 1:1000 in blocking buffer and incubated overnight at 4 °C with rocking. The blots were washed 3 x 5 minutes in phosphate buffered saline 0.1% Tween 20, then secondary antibody at a 1:15,000 dilution in blocking buffer for 1 hour at room temperature with rocking. Blots were washed again 3 x 5 minutes, then developed using SuperSignal West Chemiluminescent Substrate (Thermo Fisher, cat #34096) for 5 minutes with rocking. The membranes were protected in a plastic sleeve and imaged on an iBright CL1500 (Invitrogen Thermo Fisher).

### Quantitative Polymerase Chain Reaction (qPCR)

For validation of CRISPR knockouts, RNA was extracted from CRISPR clones using the RNeasy Mini Kit (Qiagen, cat #74104). cDNA library generation was performed using a High-Capacity cDNA Reverse Transcription Kit (Thermo Fisher Scientific, cat #4368814). Primers for VDAC1, 2, and 3 were selected from PrimerBank^50^ and obtained from Genewiz (Waltham, MA). qPCR was performed using an Applied Biosystems PowerUp SYBR Green Master Mix (Thermo Fisher Scientific, cat #A25742), with relative expression determined by ΔΔC_T_.

### BH3 Profiling

The protocol for BH3 profiling of cell lines and tissues has been described elsewhere^51^. Briefly, 2X peptide solutions were prepared (in the presence or absence of 2X inhibitor compounds or controls) with 2X digitonin (0.002%) in Mannitol experimental buffer (MEB) (10 mM HEPES pH 7.5, 150 mM Mannitol, 50 mM KCl, 0.02 mM EGTA, 0.02 mM EDTA, 0.1% BSA, 5 mM Succinate), dispense at 25 µL per well, and stored at -80 °C until use. Peptides were obtained from Biosynth (Gardner, MA); peptide sequences were BIM (Ac-MRPEIWIAQELRRIGDEFNA-NH2) and positive control DFNA5 (Ac-LQIIPTLCHLLRALSDDGVS-NH2).

Cultured cell lines of interest were harvested and centrifuged at 300 x g for 5 minutes. The pellets were resuspended in MEB and seeded at a concentration of 10,000 cells/well. Prepared profiling plates were thawed and loaded with 25 µL cell suspension, bringing the peptide/inhibitor mixture to a 1X concentration. The cells were incubated at 28 °C in a dry incubator for 1 hour before fixation with 15 µL/well of an 8% paraformaldehyde (PFA) solution in PBS. The cells were fixed for 10 minutes before neutralizing the PFA with 35 µL/well N2 neutralizing buffer (1.7 M Tris base, 1.25 M Glycine, pH 9.1, sterile filtered). A staining solution was prepared in 10X intracellular staining buffer () containing AF647-conjugated cytochrome *c* antibody (Biolegend, cat #612310) diluted 1:1000 and DAPI staining solution (Abcam, cat #ab228549) diluted 1:100. 10 µL/well staining solution was added and incubated to stain for 24 hours at 4 °C with rocking. Plates were recorded with an AttuneNxT flow cytometer (Thermo Fisher Scientific).

### Recombinant BAX Production

Recombinant BAX was produced according to a published protocol^23^. Briefly, the expression plasmid pTYB1-BAX was transformed into One Shot BL21 Star (DE3) chemically competent cells (Thermo Fisher Scientific, cat #C601003). Competent cells were cultured until an OD_600_ of 2-3 was reached, then expression of the construct, in which BAX is fused to an intein-chitin binding domain (CBD) was induced with IPTG. Cells were pelleted and lysed by sonication, then the lysate was loaded onto a chitin column to capture the BAX-intein-CBD fusion. To elute the BAX protein, DTT was added to induce cleavage of the intein domain. The eluted BAX was fractionated by FPLC using a SEC650 size-exclusion column (Bio-Rad, cat #7801650), and the fractions were analyzed by Coomassie and Western Blot analysis. Selected fractions were pooled, concentrated using a centrifugal filter, and snap-frozen for storage at -80°C.

### Fluorescence polarization Ligand Assay for Monitoring BAX Early-activation (FLAMBE)

The protocol for generation and analysis of FLAMBE data has been described elsewhere^52,53^. Briefly, working stocks of recombinant BAX, 5-TAMRA-labeled BAK-BH3 peptide (AnaSpec, cat #AS-64590), and serially diluted candidate inhibitor compounds were prepared at a working 4× concentration in 0.5X PBS. The assay was performed in a 100 μL volume per well in an opaque black polystyrene 96-well plates (Corning, cat #3915). The final concentration of BAK^TAMRA^ was 50 nM, and the final concentration of recombinant BAX was either 65 nM or 90 nM as optimized per recombinant protein preparation and indicated in the figure legends. Compound titrations were generated in-well via serial dilutions followed first by addition of BAX and finally BAK^TAMRA^ peptide before immediately being subjected to spectrometry. Fluorescence polarization was measured for each well at least once per minute for 60 minutes using a BioTek Synergy H1 plate reader (Agilent, model #8040562) equipped with a red polarization filter (Agilent/BioTek, cat #8040562). Plate reader settings were as follows: Protocol, [25 °C, 2 s double orbital shake, read]; Excitation, 530/25 nm; Excitation, 590/35 nm; Mirror, 570 nm; Gain (voltage), 50; Optics position, top; Read height, 7.0 mm. Data are the mean of technical replicates and are representative of repeated and reproduced assays.

Polarization in milli-Polarization (mP) units was calculated by the Gen5 plate reader software. For each tested drug or control dose, two parameters were extracted from the polarization time-course data: 1) endpoint fluorescence polarization (EP), or the polarization in mP at 60 minutes; and 2) time at max fluorescence polarization (Tmax), or the timepoint at which the maximum fluorescence polarization occurred. EP and Tmax values for each sample were normalized to values for BAK^TAMRA^ alone and BAX+BAK^TAMRA^ controls, as 0 and 1, respectively. for each condition was normalized to the EP for the BAK^TAMRA^ alone and no-inhibitor BAX+BAK^TAMRA^ conditions on a 0 to 1 scale. Tmax of the BAK^TAMRA^ control or any sample exhibiting no binding kinetics during the assay was set to 0 to avoid misidentification due to noise. As a single-parameter metric, the shift of each tested condition relative to the BAX+BAK^TAMRA^ control was determined using a distance formula (square root of the sum of squared differences of EP and Tmax).

### Microscale Thermophoresis (MST)

Recombinant BAX was labeled for microscale thermophoresis using a RED-NHS 2nd Generation Protein Labeling Kit (Nanotemper, cat #MO-L011) according to manufacturer instructions, using a 3-fold excess of dye during labeling. A280 and A650 values were measured using a Nanodrop and used to calculate the degree of labeling as 1.496. Aliquots of labeled rBAX were snap frozen and stored at -80°C. MST buffer was prepared by supplementing gel filtration buffer with TCEP and PEG8000 to final concentrations of 1 mM and 0.1% w/v, respectively. On the day of MST experiments, fresh digitonin (Cayman, cat #14952) was dissolved from powder and added to MST buffer to a final concentration of 0.1% w/v. Fluorescent labeled rBAX was thawed on ice, centrifuged for 5 minutes at 15,000 x g, 4°C to remove aggregates, and diluted to 10 nM in MST buffer + 0.1% digitonin. Candidate inhibitors were diluted in MST buffer + 0.1% digitonin to a high dose of 5 mM, then serially diluted in MST buffer + 0.1% digitonin with DMSO added to maintain a constant concentration of DMSO across the dilution series. rBAX was mixed with diluted compounds in equal volume, yielding a final BAX concentration of 5 nM, and the mixture was added to Monolith Premium Capillaries (Nanotemper, cat #MO-K025) for MST using a Monolith NT.115 Pico at the Harvard Medical School Center for Macromolecular Interactions. Monolith software was used to exclude samples with evidence of aggregation during thermophoresis and to calculate Fnorm values for the remaining samples.

### TMRE and Annexin V Assays

Cell lines of interest were seeded in a black-walled 96-well plate (Corning cat no. 3603) at a concentration of 1x10^4^ cells per well in 100 µL standard culture media and allowed to attach for 24 hours. On the day of treatment, solutions of candidate inhibitors and vehicle controls were prepared in standard culture media. A 6-point dose response of lethal drugs was prepared in the same manner, with a DMSO vehicle control and FCCP (Abcam, cat #ab120081) positive control. Culture media was removed from the cells and replaced with 50 µL of fresh culture media. 25 µL of 4X inhibitor or vehicle control was applied, followed by 25 µL 4X lethal drug and positive/negative controls, bringing the final concentration to 1X. Drug-containing media was left on the cells for the specified treatment time. 30 minutes prior to the end of treatment, TMRE dye diluted in media was added to the treatment media for a final concentration of 50 nM TMRE.

At the end of the treatment, the treatment media was transferred to a plate to retain any dead or dying cells that had detached, and the cells were washed with PBS and trypsinized. The trypsinized cells were then quenched with the treatment media and transferred to a U-bottom 96-well plate (Corning cat no. 351177). A 10X Annexin V staining solution (1:200 Annexin V in 100 mM HEPES (pH 7.4), 40 mM KCl, 1.4 M NaCl, 7.5 mM MgCl_2_, and 25 mM CaCl_2_) was then applied and incubated for 15 minutes on ice in the absence of light. Stained cells were analyzed using an Attune NxT (Thermo Fisher Scientific).

### Live Cell Time-Lapse Imaging

For live cell imaging, cells were seeded in black wall, clear bottom 96-well plates (Corning, cat #3603) at a density of 10,000 cells/well. The day after seeding, cells were stained for 30 minutes with Hoechst 33342 (Abcam, cat #ab228551) diluted in staining media to a concentration of 2 µM. During staining, drug dilutions were prepared in culture media for addition to the plates. Stained cells were washed with PBS, then prepared treatment media was applied to cells. Treatment media included SYTOX Green dead cell dye (Thermo Fisher Scientific, cat #S7020) at a final concentration of 20 nM. Imaging was carried out using a CellCyte X live cell imager (Cytena) housed in a tissue culture incubator in standard culture conditions, acquiring images every 4 hours. Images from the first timepoint acquired (t=0) were discarded due to condensation causing out-of-focus images at this timepoint, as were any other wells where the instrument failed to acquire the correct focus. CellCyte software was used to quantify total (Hoechst+) and dead (SYTOX Green+) objects in each acquired image. Object recognition parameters were optimized for the cell lines and stains utilized. Object counts were used to calculate a Lethal Fraction (LF) metric for each timepoint and condition, representing the fraction of total cells considered dead using the dead cell dye.

