## Supplementary Figures and Figure Legends for "A peptide-based screen for cell death inhibitors identifies the cytoprotective compound CDL36"

Figure S1

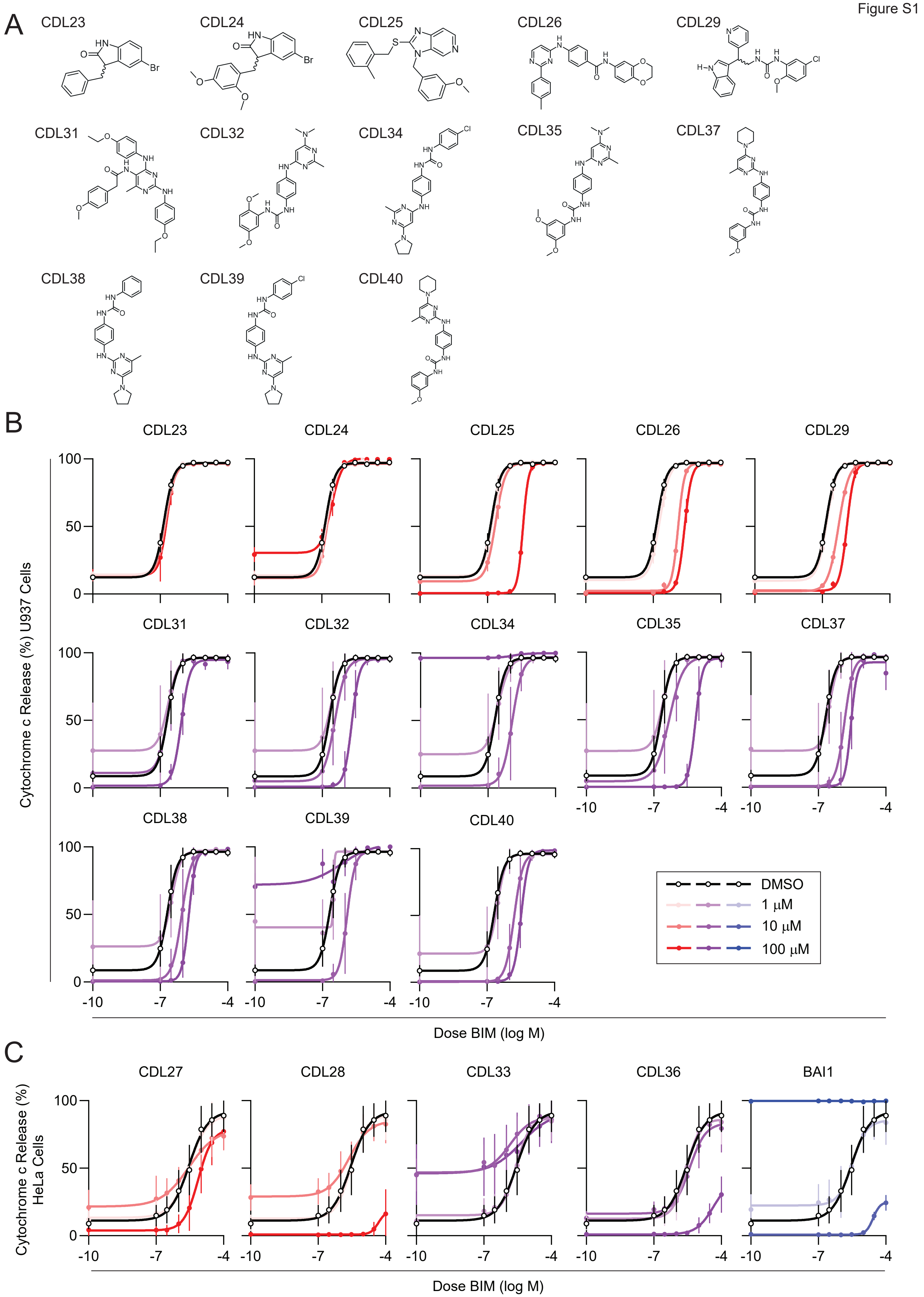
A
CDL23
CDL24
CDL25
CDL26
CDL29
CDL31
CDL32
CDL34
CDL35
CDL37
CDL38
CDL39
CDL40
B
Cytochrome c Release (%) U937 Cells
Dose BIM (log M)
DMSO
1 μM
10 μM
100 μM
C
CDL27
CDL28
CDL33
CDL36
BAI1
Cytochrome c Release (%) HeLa Cells
Dose BIM (log M)

**Figure S1. MOMP inhibition by additional candidate BAX/BAK inhibitors and in additional cell lines, related to Figure 1**

A. Small molecule structures. Structures of compounds selected for further follow-up are depicted in Figure 2.

B. Activity of selected hit compounds in U937 cells co-treated with the indicated compound doses and a range of BIM peptide doses. Data represent the mean and SD of cytochrome c signal. n = 3 biological replicates.

C. Activity of selected hit compounds and reported BAX inhibitor BAI1 in HeLa cells co-treated with the indicated compound doses and a range of BIM peptide doses. Data represent the mean and SD of cytochrome c signal. n = 3 biological replicates.

Figure S2

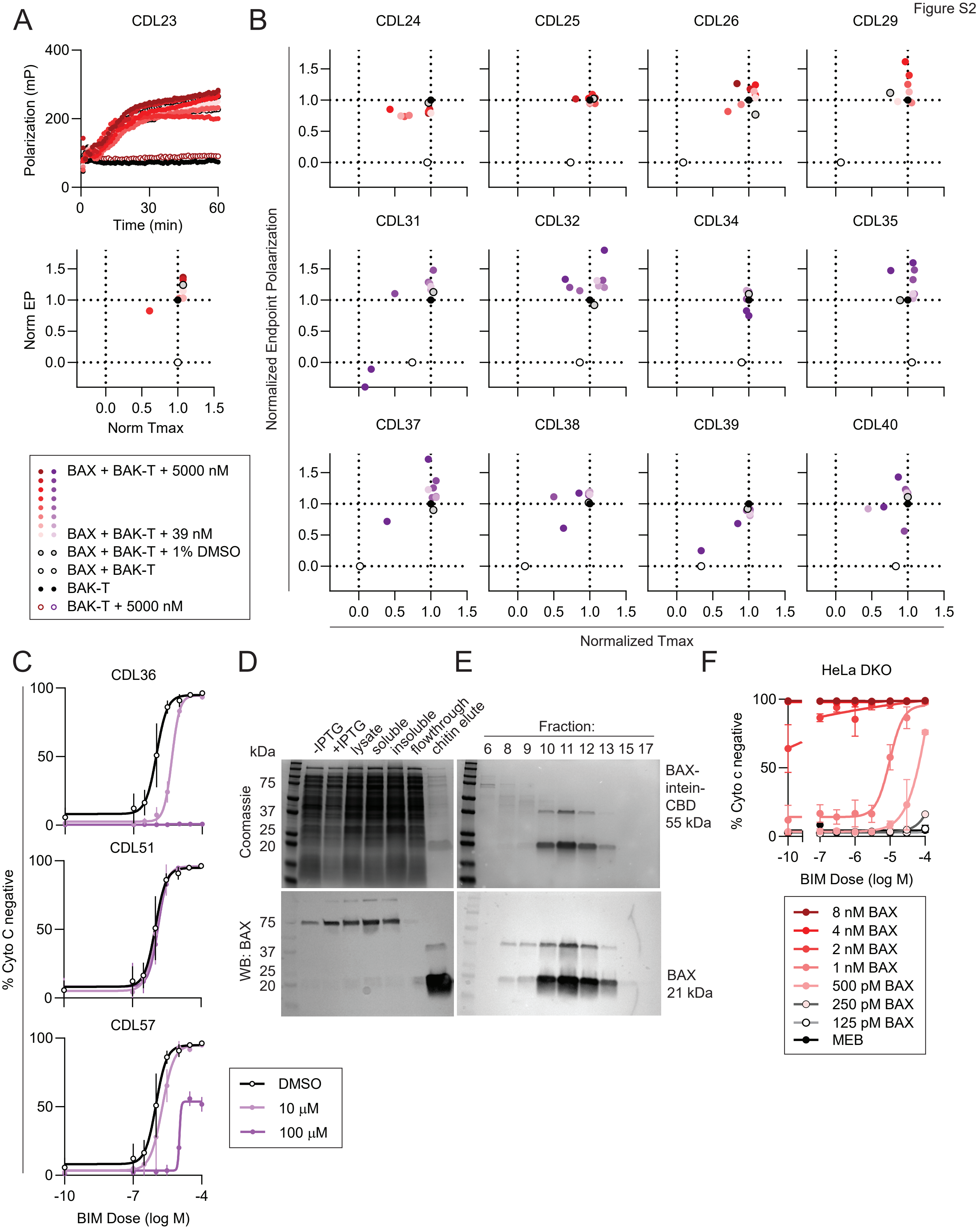

A
CDL23
Polarization (mP)
Time (min)
Norm EP
Norm Tmax
BAX + BAK-T + 5000 nM
BAX + BAK-T + 39 nM
BAX + BAK-T + 1% DMSO
BAX + BAK-T
BAK-T
BAK-T + 5000 nM
B
CDL24
CDL25
CDL26
CDL29
CDL31
CDL32
CDL34
CDL35
CDL37
CDL38
CDL39
CDL40
Normalized Endpoint Polaarization
Normalized Tmax
C
CDL36
CDL51
CDL57
% Cyto C negative
BIM Dose (log M)
DMSO
10 μM
100 μM
D
kDa
-IPTG
+IPTG
lysate
soluble
insoluble
flowthrough
chitin elute
Coomassie
WB: BAX
E
Fraction:
6 8 9 10 11 12 13 15 17
BAX-intein-CBD 55 kDa
BAX 21 kDa
F
HeLa DKO
% Cyto c negative
BIM Dose (log M)
8 nM BAX
4 nM BAX
2 nM BAX
1 nM BAX
500 pM BAX
250 pM BAX
125 pM BAX
MEB

**Figure S2. Additional FLAMBE data and generation and characterization of recombinant BAX, related to Figure 2**

A. FLAMBE assay measuring fluorescence polarization of TAMRA-labeled BAK BH3 peptide. Raw polarization data over time (top) and extracted kinetic parameters across doses (bottom) are shown for BAK-TAMRA and recombinant BAX in the presence of CDL23 or controls. Data represent average of two technical replicates.

B. Kinetic parameters extracted from FLAMBE polarization data for the indicated candidate compound or control Data represent average of two technical replicates.

C. Coomassie total protein stain (top) and Western blot for BAX protein (bottom) at the indicated stages of the recombinant BAX production and purification process.

D. Coomassie total protein stain (top) and Western blot for BAX protein (bottom) in the indicated high performance liquid chromatography (HPLC) fractions. Fractions 9-13 were subsequently pooled for concentration, characterization, and storage of recombinant BAX protein.

E. Activity of recombinant BAX in digitonin-permeabilized HeLa BAX/BAK double knockout cells as measured by BH3 profiling. HeLa cells were co-treated with a range of BIM peptide doses and the indicated concentrations of recombinant BAX. Data represent the mean and SD of cytochrome c signal. n = 3 biological replicates.

Figure S3

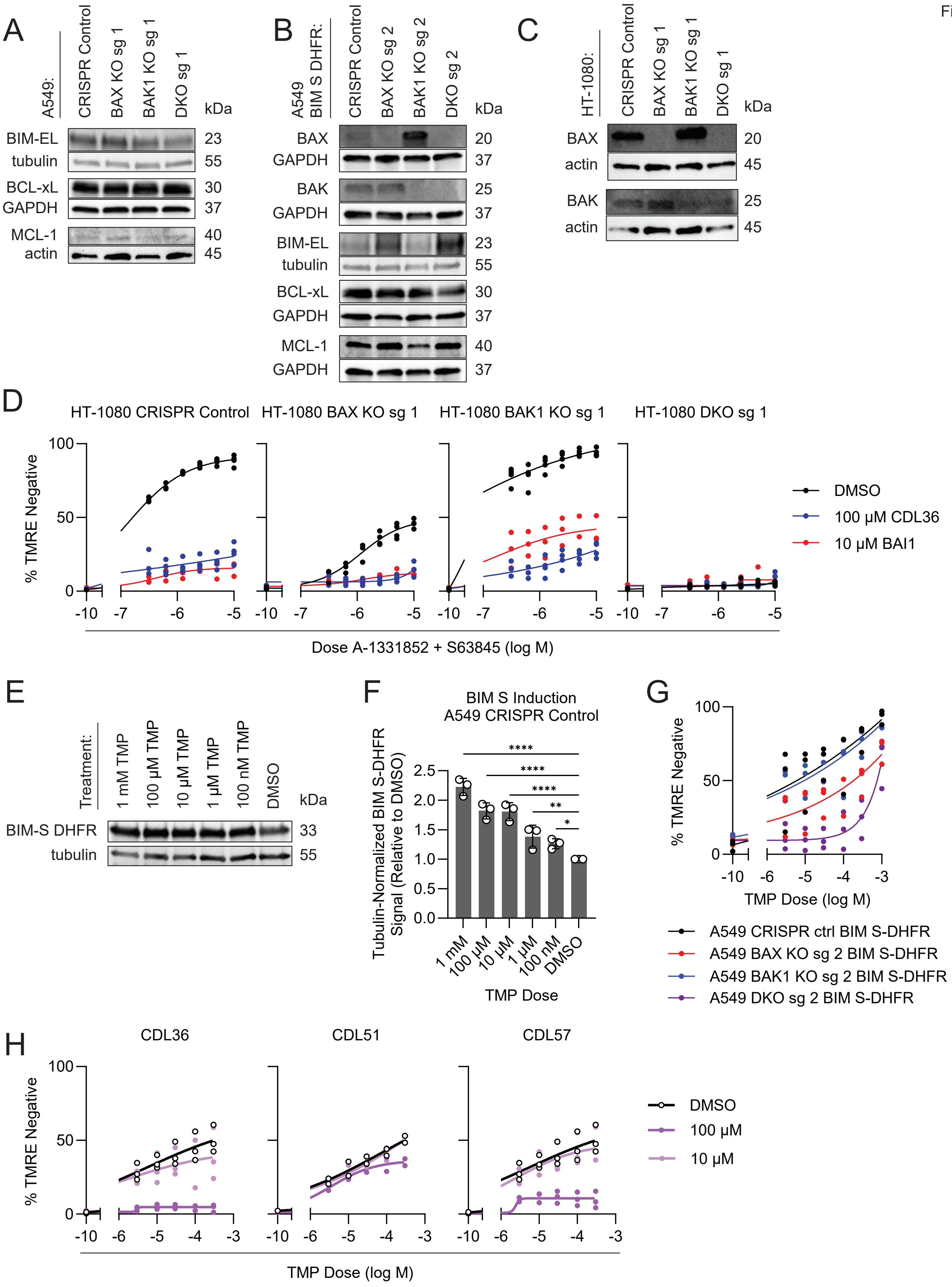

A
A549:
CRISPR Control
BAX KO sg 1
BAK1 KO sg 1
DKO sg 1
kDa
BIM-EL 23
tubulin 55
BCL-xL 30
GAPDH 37
MCL-1 40
actin 45
B
A549 BIM S DHFR:
CRISPR Control
BAX KO sg 2
BAK1 KO sg 2
DKO sg 2
kDa
BAX 20
GAPDH 37
BAK 25
GAPDH 37
BIM-EL 23
tubulin 55
BCL-xL 30
GAPDH 37
MCL-1 40
GAPDH 37
C
HT-1080:
CRISPR Control
BAX KO sg 1
BAK1 KO sg 1
DKO sg 1
kDa
BAX 20
actin 45
BAK 25
actin 45
D
HT-1080 CRISPR Control
HT-1080 BAX KO sg 1
HT-1080 BAK1 KO sg 1
HT-1080 DKO sg 1
% TMRE Negative
Dose A-1331852 + S63845 (log M)
DMSO
100 µM CDL36
10 µM BAI1
E
Treatment:
1 mM TMP
100 µM TMP
10 µM TMP
1 µM TMP
100 nM TMP
DMSO
kDa
BIM-S DHFR 33
tubulin 55
F
BIM S Induction
A549 CRISPR Control
Tubulin-Normalized BIM S-DHFR Signal (Relative to DMSO)
****
****
****
**
*
1 mM
100 µM
10 µM
1 µM
100 nM
DMSO
TMP Dose
G
% TMRE Negative
TMP Dose (log M)
A549 CRISPR ctrl BIM S-DHFR
A549 BAX KO sg 2 BIM S-DHFR
A549 BAK1 KO sg 2 BIM S-DHFR
A549 DKO sg 2 BIM S-DHFR
H
CDL36
CDL51
CDL57
% TMRE Negative
TMP Dose (log M)
DMSO
100 µM
10 µM

**Figure S3. Characterization of BIM-DHFR system, CRISPR knockouts, and inhibitor activity in an additional cell line, related to Figure 3**

A. Western blot analysis of additional BCL-2 family proteins in knockout and control A549 cells.

B. Western blot analysis of BAK/BAK KO and control lines generated with sgRNA #2.

C. Western blot analysis of knockout and control HT-1080 fibrosarcoma cells generated by CRISPR-mediated disruption of the *BAX* and *BAK1* loci.

D. CDL36 and BAI1 activity in HT-1080 cells as measured by TMRE signal loss as an indicator of MOMP. Cells were co-treated for 1 h with a range of doses of the combined BCL-xL inhibitor A-133852 and MCL-1 inhibitor S63845 and the indicated inhibitor dose or vehicle control.

E. Western blot analysis of TMP-dependent induction of BIM expression in DHFR fusion-expressing A549 cells. Image is representative of 3 biological replicates.

F. Quantification of H. Error bars represent mean +/- SD. P Values by one-way ANOVA with Holm-Šidák multiple comparisons test: ****, $P<0.0001$; **, $P<0.01$; *, $P<0.05$

G. Apoptotic sensitivity of wild-type and knockout A549 cells as measured by TMRE signal upon treatment with a range of TMP doses. n = 3 biological replicates.

H. Activity of CDL36 and structural analogs as measured by TMRE stain for MOMP upon BIM induction. BIM expression was induced by 1 h treatment with a range of TMP doses, co-treated with the indicated compound dose or vehicle control. n = 3 biological replicates.

Figure S4

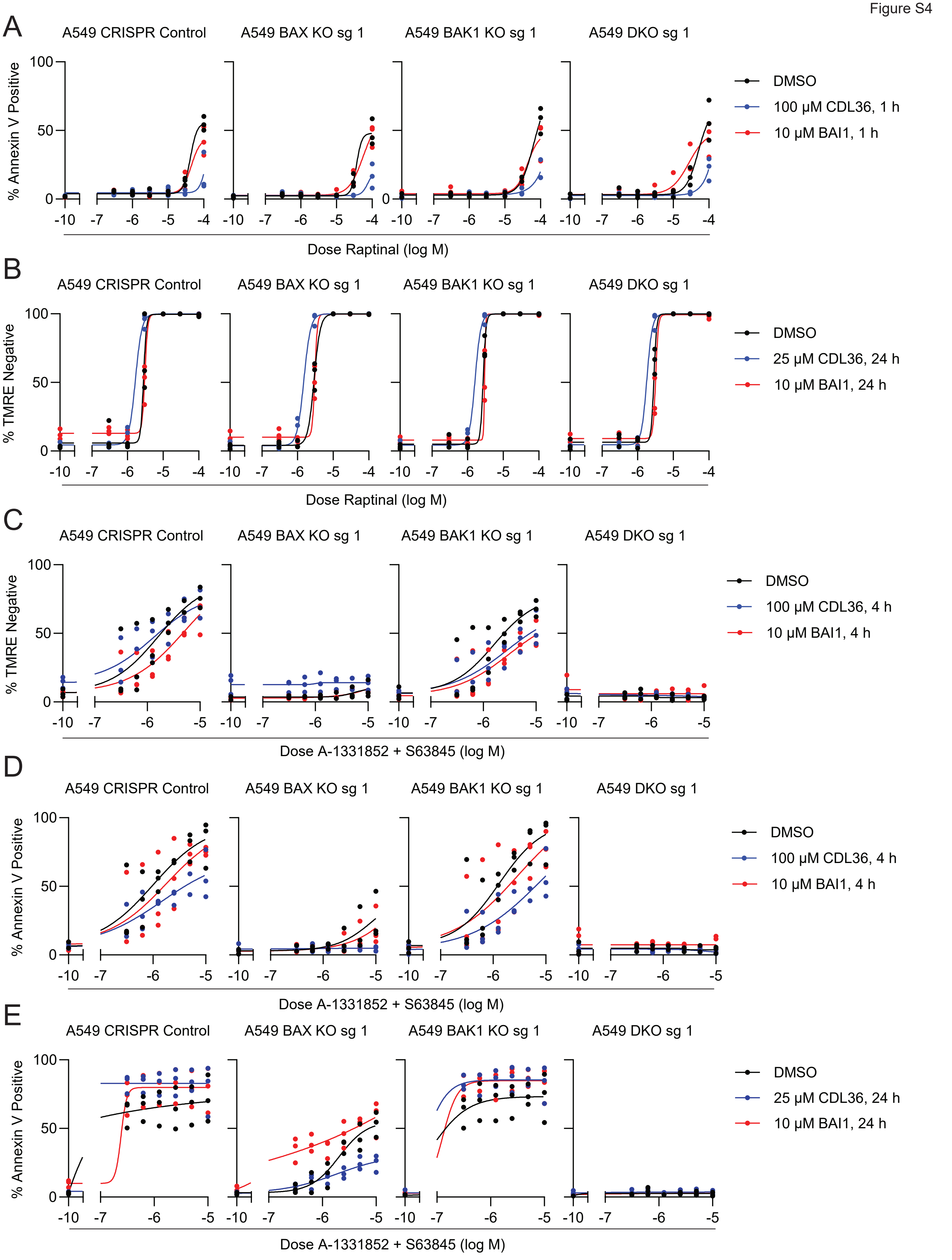

A
A549 CRISPR Control
A549 BAX KO sg 1
A549 BAK1 KO sg 1
A549 DKO sg 1
% Annexin V Positive
Dose Raptinal (log M)
DMSO
100 µM CDL36, 1 h
10 µM BAI1, 1 h
B
A549 CRISPR Control
A549 BAX KO sg 1
A549 BAK1 KO sg 1
A549 DKO sg 1
% TMRE Negative
Dose Raptinal (log M)
DMSO
25 µM CDL36, 24 h
10 µM BAI1, 24 h
C
A549 CRISPR Control
A549 BAX KO sg 1
A549 BAK1 KO sg 1
A549 DKO sg 1
% TMRE Negative
Dose A-1331852 + S63845 (log M)
DMSO
100 µM CDL36, 4 h
10 µM BAI1, 4 h
D
A549 CRISPR Control
A549 BAX KO sg 1
A549 BAK1 KO sg 1
A549 DKO sg 1
% Annexin V Positive
Dose A-1331852 + S63845 (log M)
DMSO
100 µM CDL36, 4 h
10 µM BAI1, 4 h
E
A549 CRISPR Control
A549 BAX KO sg 1
A549 BAK1 KO sg 1
A549 DKO sg 1
% Annexin V Positive
Dose A-1331852 + S63845 (log M)
DMSO
25 µM CDL36, 24 h
10 µM BAI1, 24 h

**Figure S4. Analysis of additional timepoints and stains for modulation of apoptotic signaling, related to Figure 4**

A. CDL36 and BAI1 activity as measured by Annexin V staining for PS externalization as an early indicator of apoptotic cell death. Cells were co-treated for 4 h with a range of doses of the BAX/BAK-independent apoptosis inducer raptinal and the indicated inhibitor dose or vehicle control. n = 3 biological replicates.

B. CDL36 and BAI1 activity as measured by TMRE signal loss as an indicator of MOMP. Cells were co-treated for 24 h with a range of doses of raptinal and the indicated inhibitor dose or vehicle control. n = 3 biological replicates.

C-D. CDL36 and BAI1 activity as measured by TMRE signal loss (C) or Annexin V staining (D). Cells were co-treated for 4 h with a range of doses of combined A-1331852, S63845, and the indicated inhibitor dose or vehicle control. n = 3 biological replicates.

E. CDL36 and BAI1 activity as measured by Annexin V staining. Cells were co-treated for 24 h with a range of doses of combined A-1331852, S63845, and the indicated inhibitor dose or vehicle control. n = 3 biological replicates.

Figure S5

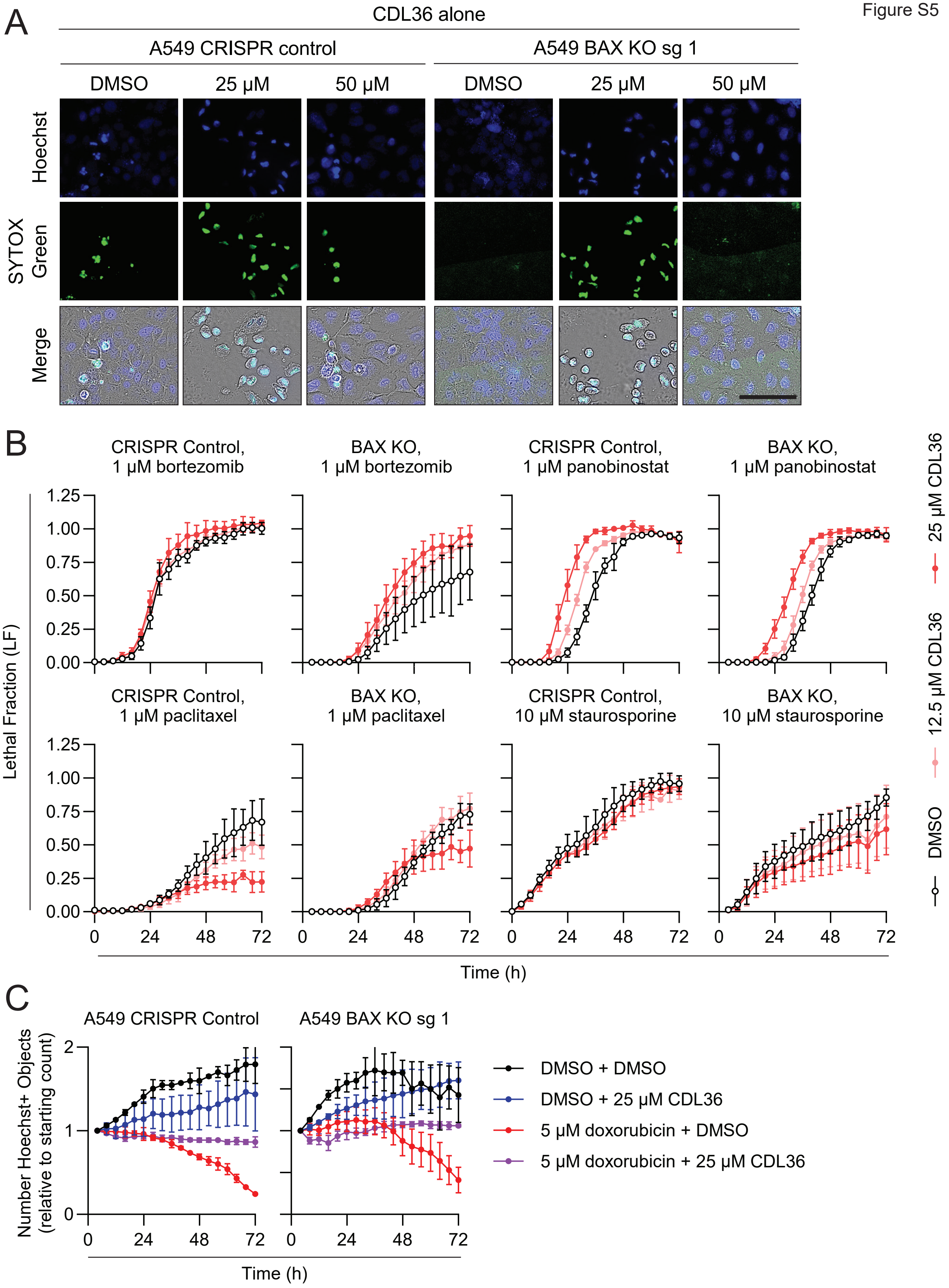

A
CDL36 alone
A549 CRISPR control
A549 BAX KO sg 1
DMSO
25 μM
50 μM
Hoechst
SYTOX Green
Merge
B
CRISPR Control, 1 μM bortezomib
BAX KO, 1 μM bortezomib
CRISPR Control, 1 μM panobinostat
BAX KO, 1 μM panobinostat
CRISPR Control, 1 μM paclitaxel
BAX KO, 1 μM paclitaxel
CRISPR Control, 10 μM staurosporine
BAX KO, 10 μM staurosporine
Lethal Fraction (LF)
Time (h)
25 μM CDL36
12.5 μM CDL36
DMSO
C
A549 CRISPR Control
A549 BAX KO sg 1
Number Hoechst+ Objects (relative to starting count)
Time (h)
DMSO + DMSO
DMSO + 25 μM CDL36
5 μM doxorubicin + DMSO
5 μM doxorubicin + 25 μM CDL36

**Figure S5. Additional timelapse quantification and representative images, related to Figure 5**

A. Representative images depicting total (Hoechst+) or dead (SYTOX Green+) cells after 48 hours of treatment with the indicated doses of CDL36. Scale bar = 100 μm.

B. LF timecourses comparing death over time in response to bortezomib, paclitaxel, panobinostat, and staurosporine in the presence or absence of the indicated CDL36 doses. Data represent the mean and SD of 3-4 biological replicates.

C. Total (Hoechst+) cell counts over time in the indicated conditions, normalized to the cell count at the first quantified timepoint. Data represent the mean and SD of 2-4 biological replicates.
