## Supplementary material for "A peptide-based screen for cell death inhibitors identifies the cytoprotective compound CDL36": Table S1

**Table S1. Description of Small Molecule Libraries**

Asterisks indicate partially screened libraries

| **Library Name** | **Library Description (from iccb.med.harvard.edu)** | **Number of Compounds in Library** | **Information Link** |
| --- | --- | --- | --- |
| NINDS Custom Collection 1 | Three quarters of the compounds in this collection are FDA-approved and the rest – which include some controlled substances such as cannabinoids – are known to influence brain activity. | 1040 | https://iccb.med.harvard.edu/ninds-custom-collection-2 |
| Gray Kinase Inhibitor Focused Library | This is a kinase inhibitor-focused library that comprises validated and suspected ATP-site kinase inhibitors that target active and inactive kinase conformations. | 187 | https://iccb.med.harvard.edu/gray-kinase-inhibitor-focused-library |
| Microsource 1 - US Drug Collection | According to MicroSource, the compounds in this collection “have reached clinical trial stages in the USA. Each compound has been assigned USAN or USP status and is included in the USP Dictionary (U.S. Pharmacopeia, 2005), the authorized list of established names for drugs in the USA”. | 1040 | https://iccb.med.harvard.edu/microsource-1-us-drug-collection |
| LOPAC 1 | This is a collection of 1280 pharmacologically-active compounds. | 1280* | https://iccb.med.harvard.edu/lopac1 |
| MSDiscovery 1 | This is a collection of compounds newly available at MicroSource (part of their Pharmakon 1600 library) and not previously represented in ICCB-L Known Bioactives libraries. | 270 | https://iccb.med.harvard.edu/msdiscovery-1 |
| LINCS1 - Kinase Inhibitors | This is a collection of characterized kinase inhibitors purchased from a variety of vendors. | 88 | https://iccb.med.harvard.edu/lincs1-kinase-inhibitors |
| Biomol ICCB-L Known Bioactives 2012 | This library includes many classes of compounds including ion channel blockers, GPCR ligands, second messenger modulators, nuclear receptor ligands, actin and tubulin ligands, kinase inhibitors, protease inhibitors, gene regulation agents, lipid biosynthesis inhibitors, as well as other well-characterized compounds that perturb cell pathways. | 480 | https://iccb.med.harvard.edu/biomol-iccb-l-known-bioactives-2012 |
| ChemBridge Focused G-Protein Coupled Receptor Library | This collection contains 250 compounds selected from ChemBridge’s larger GPCR Library of over 13,000 novel compounds. This focused collection has 11 different scaffolds, with each scaffold having 19-24 compounds. | 250 | https://iccb.med.harvard.edu/chembridge-focused-g-protein-coupled-receptor-library |
| ChemBridge Focused Ion Channel Core | This collection contains 250 compounds selected from ChemBridge’s larger IONCore Library, which is a computationally selected library of more than 4,000 small molecules selected from ChemBridge’s CORE Library. Compounds were selected using pharmacophores generated from known ion channel actives. There are 29 target-type combinations covered in this library by 21 different scaffolds, with most scaffolds having a minimum of 5 compounds. | 250 | https://iccb.med.harvard.edu/chembridge-focused-ion-channel-core |
| ChemBridge Focused Nuclear Hormone Receptor-Based Core | This collection contains 250 compounds selected from ChemBridge’s larger Nuclear Hormone Receptor-Biased Core (NHRCore) Library, which is a computationally selected library of more than 1,200 small molecules selected from ChemBridge’s CORE Library. Compounds were selected using pharmacophores generated from known nuclear hormone receptor actives. There are 10 targets (androgen receptor, estrogen receptor, estrogen receptor modulator, farnesoid X receptor, glucocorticoid receptor, liver X receptor, PPAR, progesterone receptor, RAR, and vitamin D receptor) covered in this library by 16 different scaffolds, with each scaffold having a minimum of 5 compounds. | 250 | https://iccb.med.harvard.edu/chembridge-focused-nuclear-hormone-receptor-based-core |
| ChemBridge Focused Kinase-Based Core | This collection contains 250 compounds selected from ChemBridge’s larger kinase-biased KINACore Library, which is a computationally selected library of more than 6,000 small molecules selected from ChemBridge’s CORE Library. Compounds were selected using pharmacophores generated from known kinase actives or are predicted to interact with the ATP ligand site of kinases. There are 34 targets (ADK, Aurora, BRAF, CDK, CDK2, CDK5, CHK1, C-MET, CSFR1, EGFR, FGFR, GRK2, GSK, HER, IKK, IRAK, JAK, JNN, LCK, MYT-1, P38 MAP, PDGFR, PI3K, PKA, PKB, PKC, PLK1, RAF, SRC, SYK, TGFR, TIE2, TK, VEGFR) covered in this library by 16 different scaffolds, with each scaffold having a minimum of 5 targets. | 250 | https://iccb.med.harvard.edu/chembridge-focused-kinase-based-core |
| LINCS 3 - Chromatin Targeting Library | This is a collection of compounds that inhibit proteins characterized to interact with and/or impact chromatin structure and function.  Targets include histone deacetylases, DNA methyltransferases, polyADP ribose polymerases, histone acetyl transferases, sirtuins, histone lysine methyltransferases, histone lysine demethylases, bromodomains, methyl mark readers, HIF-1alpha prolyl hydroxylase-2, telomerase, and RNA polymerase I. | 41 | https://iccb.med.harvard.edu/lincs3-chromatin-targeting-library |
| ChemDiv 7 | This library represents ChemDiv’s Targeted Diversity Library, a set of ~50,000 drug-like compounds built around 2,500 diverse chemical scaffolds. | 49128* | https://iccb.med.harvard.edu/chemdiv-7 |
| eMolecules 2014 | This library contains 248 compounds that were selected to expand the number of FDA-approved and clinical compounds available for screening at ICCB-Longwood. Small molecules included in this library were identified via text mining from a variety of sources including ChEMBL, ClinicalTrials and DrugBank by chemists at eMolecules, and then compounds already in other ICCB-Longwood Known Bioactives Collection libraries were eliminated. | 248 | https://iccb.med.harvard.edu/emolecules |
| Cayman Biolipid 1 | This library includes prostaglandins, thromboxanes, cannabinoids, D-myo–inositol-phosphates, phosphatidylinositol-phosphates, sphingolipids, inhibitors, receptor agonists and antagonists, ceramide derivatives, and several other complex polyunsaturated fatty acids. | 831 | https://iccb.med.harvard.edu/cayman-biolipid-1 |
| LINCS 4 - Kinase Inhibitor Library | This is a collection of characterized kinase inhibitors purchased from a variety of vendors. | 169* | https://iccb.med.harvard.edu/lincs4-kinase-inhibitor-library |
| Tocriscreen™ Mini Library 3 | This is a library of 1,120 biologically active compounds. | 1120* | https://iccb.med.harvard.edu/tocriscreen%E2%84%A2-mini-library-3 |
