## Supplementary material for "A peptide-based screen for cell death inhibitors identifies the cytoprotective compound CDL36": Table S2

**Table S2. Oligonucleotide Sequences**

| **Oligonucleotide Name** | **Description** | **Sequence** |
| --- | --- | --- |
| BAX_sg1_top | *BAX* sgRNA 1 top strand | CACCGgtttcatccaggatcgagca |
| BAX_sg1_bottom | *BAX* sgRNA 1 bottom strand | AAACtgctcgatcctggatgaaacC |
| BAX_sg2_top | *BAX* sgRNA 2 top strand | CACCGaggatcgagcagggcgaatg |
| BAX_sg2_bottom | *BAX* sgRNA 2 bottom strand | AAACcattcgccctgctcgatcctC |
| BAK1_sg1_top | *BAK1* sgRNA 1 top strand | CACCGccttacctctgcaacctagc |
| BAK1_sg1_bottom | *BAK1* sgRNA 1 bottom strand | AAACgctaggttgcagaggtaaggC |
| BAK1_sg2_top | *BAK1* sgRNA 2 top strand | CACCGggcggtaaaaaacgtagctg |
| BAK1_sg2_bottom | *BAK1* sgRNA 2 bottom strand | AAACcagctacgttttttaccgccC |
| VDAC2_sg1_top | *VDAC2* sgRNA 1 top strand | CACCGtgttaggaattttcaacgtc |
| VDAC2_sg1_bottom | *VDAC2* sgRNA 1 bottom strand | AAACgacgttgaaaattcctaacaC |
| VDAC2_sg2_top | *VDAC2* sgRNA 2 top strand | CACCGaaatacaagtggtgtgagta |
| VDAC2_sg2_bottom | *VDAC2* sgRNA 2 bottom strand | AAACtactcacaccacttgtatttC |
