## Supplementary material for "A peptide-based screen for cell death inhibitors identifies the cytoprotective compound CDL36": Table S3

**Table S3. PCR and qPCR Primer Sequences**

| **Oligonucleotide Name** | **Description** | **Sequence** |
| --- | --- | --- |
| BAX_PCR_F | *BAX* locus gDNA PCR forward primer | TAGCGTTCCCCTAGCCTCTT |
| BAX_PCR_R | *BAX* locus gDNA PCR reverse primer | GAAAATTCCCGCATCCACCG |
| BAK1_PCR_F | *BAK1* locus gDNA PCR forward primer | TGGGATATTTCTTTCCATGGGTGA |
| BAK1_PCR_R | *BAK1* locus gDNA PCR reverse primer | ATGGTTGTGACATGACAGAGGA |
| VDAC2_PCR_F | *VDAC2* locus gDNA PCR forward primer | ACAGTGCAGAAGGATACGAGT |
| VDAC2_PCR_R | *VDAC2* locus gDNA PCR reverse primer | CTTACCCTGTGTTTGGTGAGAA |
| VDAC1_qPCR_F | *VDAC1* qPCR forward primer | ACGTATGCCGATCTTGGCAAA |
| VDAC1_qPCR_R | *VDAC1* qPCR reverse primer | TCAGGCCGTACTCAGTCCATC |
| VDAC2_qPCR_F | *VDAC2* qPCR forward primer | GCTACAGGACTGGGGACTTC |
| VDAC2_qPCR_R | *VDAC2* qPCR reverse primer | AATGCCAAAACGAGTGCAGTT |
| VDAC3_qPCR_F | *VDAC3* qPCR forward primer | TCAGATGAGTTTTGACACAGCC |
| VDAC3_qPCR_R | *VDAC3* qPCR reverse primer | GAAGTCCGCAGCCTTGTAAC |
| attB1_F_Cit-tevs-DHFR | Forward primer for amplification of Cit-tevs-DHFR sequence for Gateway BP reaction | ggggacaagtttgtacaaaaaagcaggctttgaaggagatagaaccatgggatccgtgagcaagggcgagg |
| attB2_R_Cit-tevs-DHFR | Reverse primer for amplification of Cit-tevs-DHFR sequence for Gateway BP reaction | ggggaccactttgtacaagaaagctgggttcatcggcgctccaatatttcaaag |
| RC_F_BamHI_BimEL | Forward primer appending BamHI site to 5’ end of Bim EL sequence for restriction cloning | GAATTAGGATCCagcccgggcggttccatg |
| RC_R_BimEL_XhoI | Reverse primer appending XhoI site to 3’ end of Bim EL sequence for restriction cloning | GACTCACTCGAGatgccttctccataccag |
